# HIDE: Hierarchical cell-type Deconvolution

**DOI:** 10.1101/2025.01.31.634483

**Authors:** Dennis Völkl, Malte Mensching-Buhr, Thomas Sterr, Sarah Bolz, Andreas Schäfer, Nicole Seifert, Jana Tauschke, Austin Rayford, Oddbjørn Straume, Helena U. Zacharias, Sushma Nagaraja Grellscheid, Tim Beissbarth, Michael Altenbuchinger, Franziska Görtler

**Affiliations:** Computational Biology Unit, Department of Biological Sciences, University of Bergen, Thormohlensgt 55, Bergen N-5008, Norway; Institute of Theoretical Physics, University of Regensburg, 93053 Regensburg, Germany; Department of Medical Bioinformatics, University Medical Center Göttingen, 37075 Göttingen, Germany; Institute of Human Anatomy and Embryology, University of Regensburg, 93053, Germany; Peter L. Reichertz Institute for Medical Informatics of TU Braunschweig and Hannover Medical School, Hannover Medical School, Carl-Neuberg-Str. 1, 30625 Hannover, Germany; Department of Biomedicine and Centre for Cancer Biomarkers, University of Bergen, Thormohlensgt 55, Bergen N-5008, Norway; Department of Oncology and Medical Physics, Haukeland University Hospital, Haukelandsveien 22, 5021 Bergen, Norway

## Abstract

**Motivation:** Cell-type deconvolution is a computational approach to infer cellular distributions from bulk transcriptomics data. Several methods have been proposed, each with its own advantages and disadvantages. Reference based approaches make use of archetypic transcriptomic profiles representing individual cell types. Those reference profiles are ideally chosen such that the observed bulks can be reconstructed as a linear combination thereof. This strategy, however, ignores the fact that cellular populations arise through the process of cellular differentiation, which entails the gradual emergence of cell groups with diverse morphological and functional characteristics.

**Results:** Here, we propose Hierarchical cell-type Deconvolution (HIDE), a cell-type deconvolution approach which incorporates a cell hierarchy for improved performance and interpretability. This is achieved by a hierarchical procedure that preserves estimates of major cell populations while inferring their respective subpopulations. We show in simulation studies that this procedure produces more reliable and more consistent results than other state-of-the-art approaches. Finally, we provide an example application of HIDE to explore breast cancer specimens from TCGA.

**Availability:** A python implementation of HIDE is available at zenodo (doi:10.5281/zenodo.14724906).

**Supplementary information:** Supplementary material is available at *Bioinformatics* online.

## 1 Introduction

The distribution of cell types within heterogeneous bulk tissues, such as tumors, plays a pivotal role in disease progression and therapeutic response. Cell-type deconvolution has emerged as a powerful and well-established computational approach to predict these distributions. Numerous deconvolution algorithms have been developed in recent years, including BayesPrism, CIBER-SORTx, MuSiC, and DTD (Chu *et al*., 2022; Newman *et al*., 2019; Wang *et al*., 2019; Görtler *et al*., 2020). By inferring the proportions of specific cell types within complex mixtures, these methods offer critical insights into cellular interactions and their roles in both physiological and pathological processes (Abbas *et al*., 2009; Altboum et al., 2014). This includes estimating the proportions of various cell types within the tumor microenvironment and linking these compositions to clinically relevant factors such as overall survival, progression-free survival, metastasis development and therapeutic response (Junttila and de Sauvage, 2013; Wang *et al*., 2023), as well as using this information to guide treatment decisions (Li *et al*., 2016; Gentles et al., 2015; Petitprez *et al*., 2018).

Despite their vast utility, the application of cell-type deconvolution faces significant challenges (Garmire *et al*., 2024; Maden et al., 2023). First, closely related cell populations typically have highly correlated molecular profiles, complicating accurate inference of cellular distributions (Görtler *et al*., 2020). Second, estimates of rare or minor cell populations are often blurred by a high signal-to-noise ratio of a major cell type (Nguyen *et al*., 2024). Both issues are typically related, as many minor cell populations share highly co-linear expression profiles.

Cellular differentiation processes result in hierarchical relationships between major and minor cell types, typically represented as cell lineage trees (Li *et al*., 2024). Minor cell types represent subsets of their respective major categories and may differ only in the expression of a single receptor. Thus, their overall population might be small and their proportions difficult to estimate as a consequence of low signal-to-noise ratios. Moreover, their expression profiles might correlate strongly to those of related minor populations and their respective major category.

Theoretically, the proportions of minor cell types should sum to the total of their corresponding major cell type. To leverage this principle, we introduce HIDE (Hierarchical cell-type Deconvolution), a novel algorithm for cell-type deconvolution. HIDE enhances the accuracy of inferring both minor and major cell populations, and more precisely disentangles closely related cellular populations. We illustrate in simulation studies that standard approaches are prone to inconsistent results, contradicting the underlying cell-type hierarchy. Furthermore, we show that HIDE substantially improves cell-type deconvolution via a machine learning procedure which optimizes cell-type deconvolution in a hierarchy-preserving manner. Finally, we provide a comprehensive analysis of cellular compositions in breast cancer, comprising association analysis with patient outcomes and a differential composition analysis comparing breast cancer subtypes.

## 2 Methods

### 2.1 Loss-function learning for cell-type deconvolution

It is well known that the selection of phenotypic marker genes is essential for reliable cell-type deconvolution (Qiu *et al*., 2021; Fischer and Gillis, 2022). Moreover, we have shown that continuous gene weighting substantially improves results (Görtler *et al*., 2020; Görtler *et al*., 2024). These gene-weights are established using single-cell data with cell-type annotation. HIDE builds on this idea and utilizes the gene-weight learning of (Görtler *et al*., 2020) at each cell-type layer for an optimal, hierarchy preserving deconvolution. We first recapitulate the concept of loss-function learning to infer gene weights.

Let 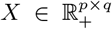 be a reference matrix with *q* cell-type specific reference profiles in their columns, and 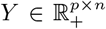 a matrix of *n* bulk gene expression profiles, both containing expression values of *p* genes in their rows. Bulk gene expression profiles can be considered as linear combinations of cell-type-specific reference profiles,

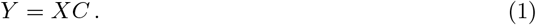

Here the columns of the cellular distribution 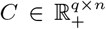 contain cellular weights/proportions corresponding to the columns (bulks) of *Y* . A single entry *C*_*ki*_ can be interpreted as semi-quantitative measurements of the number of cells of type *k* in bulk profile *i*. The cellular distribution *C* can be estimated by

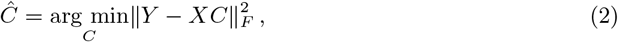

where ||.||_*F*_ is the Frobenius norm and *Ĉ* the estimate of *C* for given *X* and *Y* . Eq. (2) can be solved for each column (bulk) in *Y*, individually. In (Görtler *et al*., 2020; Schön *et al*., 2020), Eq. (2) was augmented by gene weights *g* = (*g*_1_, *g*_2_, …, *g*_*p*_)^*T*^, yielding

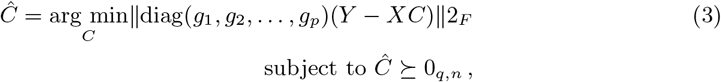

where the weights *g*_*i*_ can be chosen to improve estimates of *C*. Given bulks of known cellular compositions, the weights *g*_*i*_ can be established by minimizing

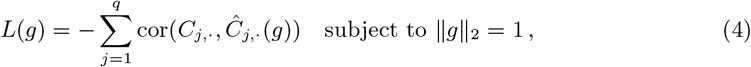

with respect to *g* = (*g*_1_, …, *g*_*p*_)^*T*^, where *C*_*j*,·_ is the *j*th row (cell type) of *C* and *Ĉ*_*j*,·_ its corresponding estimate. In other words, the Pearson’s correlation between the ground truths *C*_*j*,·_ and its estimates *Ĉ*_*j*,·_ for all cell types of interest *j* gets maximized when minimizing the loss function Eq. (4) with respect to the *g*_*i*_’s.

This strategy requires bulks *Y* of known cellular composition for gene weighing. Those are typically not available for large-scale bulk transcriptomics data. Therefore, our strategy is to build pseudo bulks from single-cell data (see Methods section about training and validation data). These pseudo bulks have known cellular composition and can hence be used for estimating the gene weights *g* as well as provide artificial mixtures for their validation.

### 2.2 HIDE algorithm

HIDE directly incorporates the cell-type hierarchy into the deconvolution procedure. We consider a cell hierarchy as a tree-like structure which relates cell types. Thus, it can be both a cell lineage tree capturing cellular differentiation processes as well as a dendrogram representing cell-type similarities. For illustration purposes, we restrict ourselves to trees consisting of three layers, only: major, minor, and sub-minor. One should note, however, that HIDE can account for arbitrary cell trees and is not limited to three layers. For the presented analysis, we retrieved the cell tree depicted in Figure 1 from DISCO.

**Figure 1:**
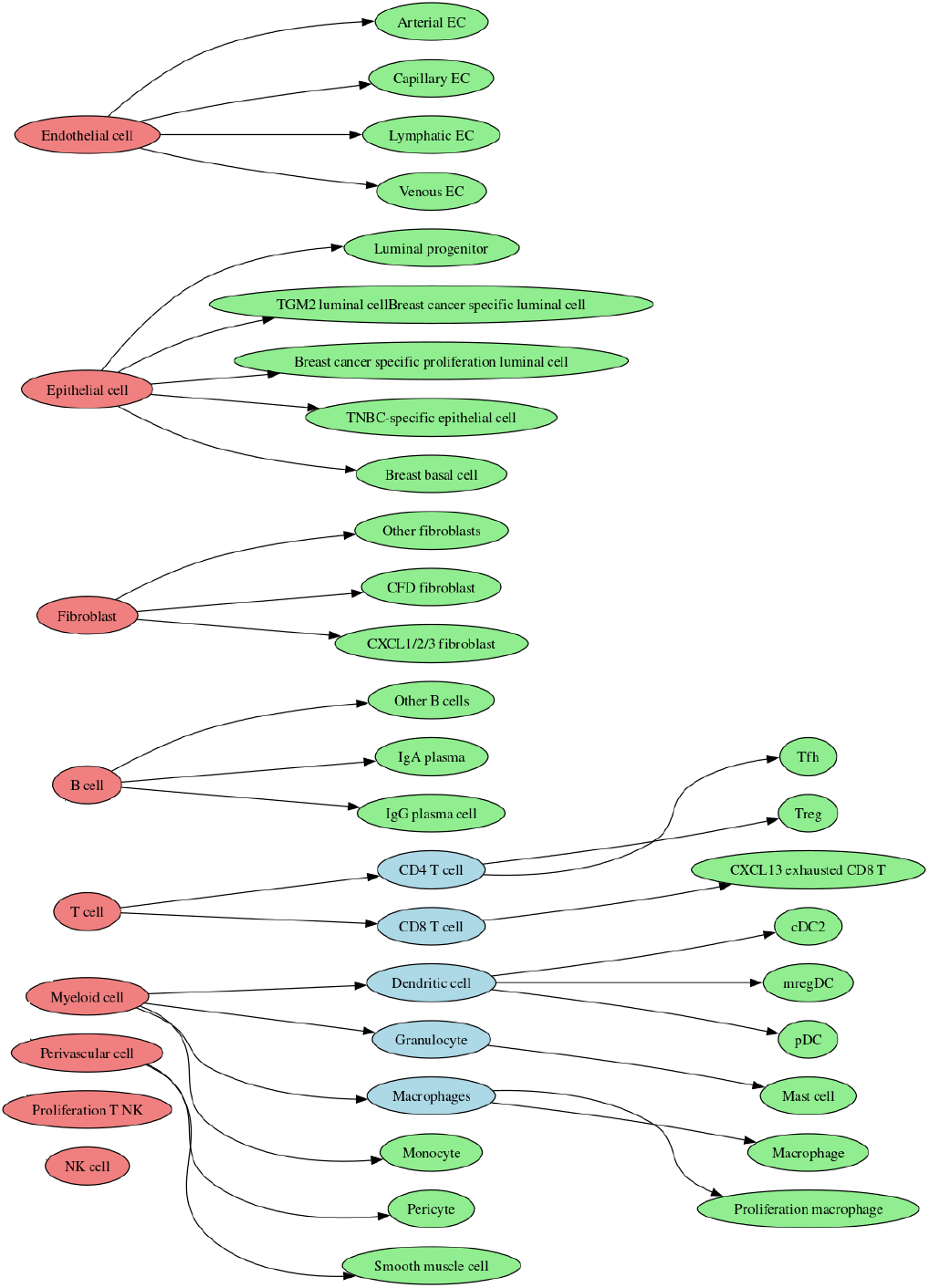
Cell type level structure for breast cancer single cell RNA-Seq dataset. Red indicates major cell type level, blue minor and green the sub-minor cell type level. When the minor cell type level is missing, the minor and major cell type are identical.

HIDE’s basic concept is to proceed through successive levels of cell-type specification, beginning with the most general cell-type level and progressing to the most specific one. At each level, the algorithm partitions the data corresponding to each parent cell type into its respective child cell types and performs deconvolution independently on the child cell types. Specifically, HIDE introduces four algorithmic components in order to incorporate cellular hierarchies into the deconvolution procedure:

#### (0) Establish a top level deconvolution

We start with the highest levels of our cell trees (the roots), corresponding to the major cell populations. We aggregate single-cell profiles representing the major cell types to generate a reference matrix *X*. Then, we learn gene weights *g* = (*g*_1_, *g*_2_, …, *g*_*p*_) via loss-function learning, where we provide the artificial training bulks *Y*, the reference matrix *X*, and respective ground-truth cellular proportions *C* aggregated to major cell types as input. The estimated cellular proportions are finally normalized to one for each bulk profile. The resulting model provides the HIDE estimates for the root cell types.

Next, we iteratively proceed through the hierarchy of the input trees performing the following three steps:

#### (1) Generate residual bulks

We first subtract the content of all but the parent cell type of interest *k* from the bulk profiles *Y* :

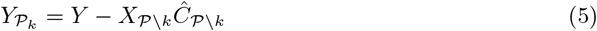

where 𝒫 is the set of all parent cell types, and 𝒫\*k* the set of all parent cell types excluding the cell type of interest *k*. The matrices *X*_𝒫\*k*_ and *Ĉ*^𝒫\*k*^ are obtained from *X* and *C* by deleting the column and row corresponding to cell type *k*, respectively. This approach ensures that the estimated proportions of subpopulations add up accurately to the total of their parent cell type, avoiding any over- or underestimation of their combined contribution. To avoid negative values in the newly created residual bulks 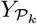, we set 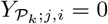 for 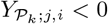.

#### (2) Establish a cell-subtype gene weighting

We establish gene weights as previously in step (0), but now use the residual bulks as input together with a reference matrix containing the subtypes of interest, only. Then, respective ground-truth cellular proportions serve as optimization target for the gene-weight learning. This step provides gene weights tailored to the deconvolution of the child cell types of interest.

#### (3) Normalize results

Finally, we normalized our results as follows.

First, one should note that the gene-weighting procedure is optimized for high Pearson’s correlation, meaning the optimization target is that estimated cellular proportions correlate well with respective ground-truth values. Thus, estimates of *C* are not optimized to agree on a global scale. We therefore incorporated a re-scaling step such that the *Ĉ*_*j*,·_ better match the *C*_*j*,·_,

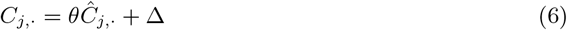

where *θ* is the slope and Δ the intercept estimated by linear regression. Negative values are set to zero. These parameters are established on the training data and remain fixed when applied to real bulk mixtures.

Second, the above procedure does not guarantee that the estimates for the parent cell coincide with the aggregated cellular proportions of its child cell types. Let 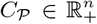 be the cellular proportions of a parent cell type and *C* ∈ ℝ^*q𝒫 ×n*^ the respective cellular proportions of its *q*𝒫 child subtypes. To enable communication between the different cellular levels, we introduce normalization factors *ξ* _𝒫_ = (*ξ* _𝒫;1_, …, *ξ* _𝒫;*n*_) ∈ ℝ^*n*^ for every bulk sample which normalize the content of all child cell types to the corresponding parent cell type:

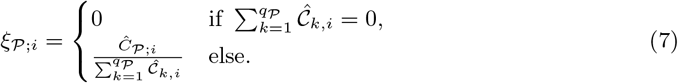

Setting *ξ*_𝒫;*i*_ to zero for 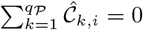 prevents division by zero. This approach helps constrain the estimates of the child cell types, preventing the algorithm from over- or underestimating their joint contribution. By definition, the algorithm is designed in a way that no contribution of a parent cell type for a given mixture also results in no contributions of the corresponding child terms.

Steps (1) to (3) are performed for each set of child cell types. This procedure is performed iteratively until models for each node in the tree are established. This is achieved when models are established for all leaf cell types.

### 2.3 Training and validation single cell dataset

For creating simulated bulk profiles with known cellular composition, we used breast cancer single-cell RNA-Seq data with provided cell type annotation from the single-cell platform DISCO (Li *et al*., 2021) (downloaded 2024-09-04). It contains 34 cell types at the finest resolution. These cells were separated into two equally sized and distributed groups for training and testing (*n* = 98, 453 vs *n* = 98, 465). Table S1 summarizes the number of cells per cell type and respective percentage values. We generated 20, 000 artificial training bulk mixtures by randomly drawing 100 cell profiles each from the training cohort (Table S1). Further, we generated 10 batches for model testing with 1, 000 samples each, following the same strategy. We then aggregated the training single-cell profiles to consensus cell-type profiles for each cell population. Finally, we retained the 5,000 genes with the highest variance for analyses (calculated across the reference matrix). This dataset provides a three-level hierarchical structure: nine major cell types, followed by 13 minor cell types (resulting from the subdivision of two major cell types), and finally 34 sub-minor-cell types. The sub-minor-cell type layer arises from the further subdivision of 11 of the 13 minor cell types. Figure S1 illustrates the underlying cell type hierarchy.

### 2.4 Competing methods

We compared HIDE to three state-of-the-art bulk deconvolution algorithms: CIBERSORTx, BayesPrism, and MuSiC (Chu *et al*., 2022; Newman *et al*., 2019; Wang *et al*., 2019). CIBER-SORTx can utilize user-specified reference profiles for deconvolution. Therefore, we employed the same reference profiles and test mixtures as used for HIDE. In contrast, BayesPrism and MuSiC operate directly on single-cell raw data. Consequently, we provided them with the single-cell profiles used to generate the reference profiles and the training data for HIDE. Both BayesPrism and MuSiC incorporate gene selection procedures; therefore, we provided all 33,538 available genes for deconvolution. Due to the computationally demanding nature of the BayesPrism algorithm, we utilized only half of the training cells. To maintain biological variability, we selected every second cell from each cell type and patient (see Table S1).

### 2.5 TCGA breast cancer analysis

Bulk transcriptomic data for breast cancer (BRCA) patients were retrieved from The Cancer Genome Atlas (TCGA) Research Network (The Cancer Genome Atlas, https://www.cancer.gov/tcga). These data encompass a total of 1,083 samples, categorized as follows: 190 triple-negative breast cancers (TNBC; ER-negative, PR-negative, HER2-negative), 82 human epidermal growth factor receptor 2-positive breast cancers (HER2+), 562 luminal A breast cancers (LumA; ER-positive, PR-positive, HER2-negative, and low Ki-67 levels), 209 luminal B breast cancers (LumB; ER-positive, PR+/-, and HER2+), and 40 breast cancer-free tissue samples (normal). To account for systematic biases introduced by differing measurement technologies, the TCGA data was subjected to gene-wise normalization using the single-cell training data as a reference. This normalization process involved calculating gene-specific re-scaling factors. These factors were obtained by dividing the mean expression of each gene in the single-cell pseudo-bulks of the training set by the corresponding mean expression observed in the TCGA control samples. For downstream analysis, we utilized survival time, days to last follow-up, and tumor stage. We further restricted the gene space to the 4,851 genes available in both the DISCO training data and the TCGA bulks. For the TCGA analysis, models were established based on this reduced gene set.

## 3 Results

### 3.1 Benchmark study

We benchmarked the performance of HIDE against three other deconvolution algorithms: BayesPrism, CIBERSORTx, and MuSiC. Since these latter algorithms, in contrast to HIDE, cannot account for the hierarchical structure between cell populations, we applied them for the inference of major and minor cell type proportions using two different strategies. The first was to aggregate sub-minor-level predictions to derive cellular proportions for the next-highest level in the hierarchy (the parent level), referred to as “summing approach” henceforth. The second approach was to perform the deconvolution for each level in the hierarchy, individually, referred to as “independent level approach”.

#### 3.1.1 Benchmark analysis

To compare performance on the level of minor and major cell populations, we then used both the summing and independent level approaches to generate predictions with BayesPrism, CIBER-SORTx, and MuSiC. This step was not necessary for HIDE, as its hierarchical top-down architecture ensures consistent results across all cell-type levels. On the level of both major and minor cell types, we found that HIDE outperformed the competing algorithms (Table 1). Considering overall performance, HIDE yielded superior compositional predictions in terms of average correlation across all cell types, with improvements over other methods ranging from 0.2 (versus CIBERSORTx on the major cell type level using the summing approach; 0.88 ± 0.003 versus 0.661 ± 0.007, respectively) up to nearly 0.4 (versus MuSiC on the major cell type level using the independent level approach; 0.88 ± 0.003 versus 0.511 ± 0.01, respectively). Notably, all four algorithms performed better on the level of major and minor cell populations than the sub-minor-cell type level. Results obtained for individual cell types are shown in Table S3 and S4 for the summing approach and in Table S5 and S6 for the individual level approach.

**Table 1:**
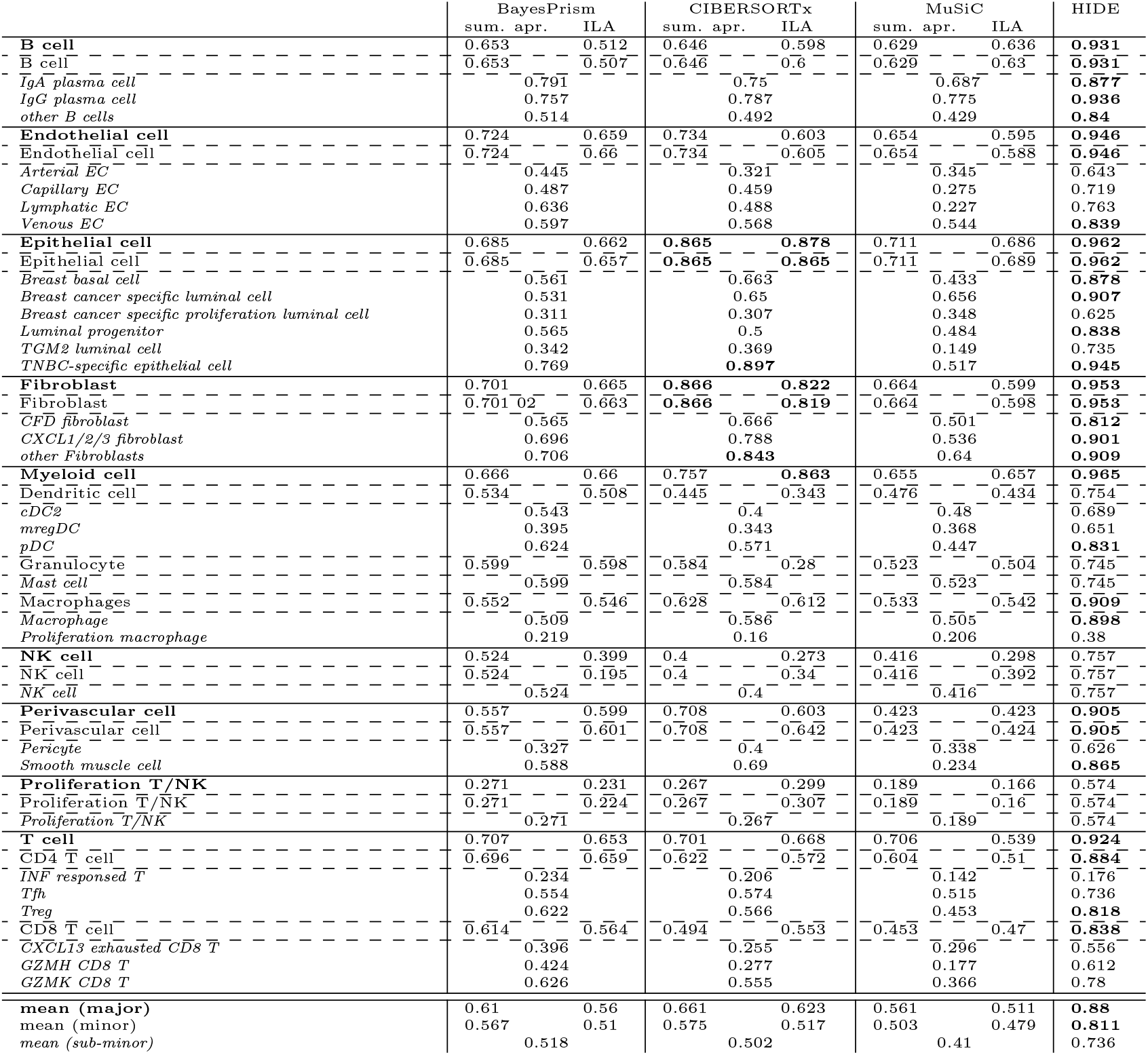
Performance comparison via Pearson’s correlation between HIDE, BayesPrism, CIBERSORTx, and MuSiC. The table is sorted by cell-type hierarchy. Bold indicates cell types on the major level, italic on sub-minor level, whereas the minor level is written in plain text. “Sum. apr.” refers to the summing approach, “ILA” to the individual level approach. Pearson’s correlations were calculated between estimated and predicted cellular proportions using the artificial test mixtures (see Methods). Results correspond to the mean correlation obtained over 10 simulation runs. Correlations above 0.8 are highlighted in bold.

An alternative benchmark analysis based on Normalized Mean Absolute Error (NMAE) values further supports our findings (Table 2, detailed results in Tables S9 - S13 in the supplement). HIDE consistently exhibited the lowest NMAE values across all levels of cellular resolution. At the individual cell type level, HIDE outperformed all competitors except for granulocytes (a minor cell type). Notably, for granulocytes and mast cells (the sole child cell type of granulocytes), BayesPrism demonstrated superior performance with an NMAE of 0.089 ± 0.01 compared to HIDE’s 0.094 ± 0.013, irrespective of whether the summing or independent level approach was employed.

**Table 2:** Performance comparison via Pearson’s correlation between summing and individual level approaches for HIDE, BayesPrism, CIBERSORTx, and MuSiC as well as mean NMAE. Bold indicates cell types on the major level, italic on sub-minor level, whereas the minor level is written in plain text. “Sum. apr.” refers to the summing approach, “ILA” to the individual level approach. Pearson’s correlations were calculated between predicted cellular proportions of summing and individual level approach. NMAE were calculated between estimated and predicted cellular proportions using the artificial test mixtures (see Methods). Results correspond to the mean correlation obtained over 10 simulation runs. Correlations above 0.9 are highlighted in bold as well as NMAE values below 0.1.

|  | BayesPrism |  | CIBERSORTx |  | MuSiC |  | HIDE |
| --- | --- | --- | --- | --- | --- | --- | --- |
|  | sum. apr. | ILA | sum. apr. | ILA | sum. apr. | ILA |  |
| **NMAE (major)** | 0.196 | 0.23 | 0.149 | 0.168 | 0.197 | 0.257 | **0.059** |
| NMAE (minor) | 0.205 | 0.242 | 0.176 | 0.214 | 0.234 | 0.266 | **0.075** |
| *NMAE (sub-minor)* | 0.196 |  | 0.186 |  | 0.263 |  | **0.086** |
| **Cor. sum. apr. vs ILA (major)** | 0.793 |  | 0.643 |  | 0.799 |  | **1** |
| Cor. sum. apr. vs ILA (minor) | 0.793 |  | 0.673 |  | 0.85 |  | **1** |

Interestingly, all three competing algorithms showed an inferior performance for the summing approach compared to the independent level approach like also found for the Pearsons’ correlation (Table 1). To further substantiate this finding, we compared the results calculated with the summing approach to those of the independent level approach using Pearson’s correlations (Table 2, detailed results in Table S7 and S8). We observed that the two approaches yielded inconsistent results, with mean correlation values as low as 0.673 ± 0.007 for CIBER-SORTx on the minor and 0.643 ± 0.009 on the major level. MuSiC achieved the most consistent values with mean correlations of 0.799 ± 0.007 and 0.85 ± 0.005 for major and minor levels, respectively. Although a correlation of 0.85 may seem relatively high, it nevertheless reveals the inherent inconsistencies between both calculation approaches.

In contrast, HIDE is by definition perfectly consistent in this analysis, as all correlation values are 1.0. This also highlights a practical advantage of HIDE; model establishment for cell types lower in the hierarchy does not affect results at higher levels, and can be stopped at any level wherein further sub-division does not yield meaningful improvements in deconvolution accuracy.

In summary, HIDE outperforms state-of-the-art methods on all levels of cellular resolution for the vast majority of cell types.

### 3.2 Ablation study

HIDE builds on the gene weight learning we proposed previously in (Görtler *et al*., 2020) and extends it by employing a cell-type hierarchy. As a next step, we therefore illustrate the effect of the additional algorithmic steps (1) to (3) (see Methods) in an ablation study, wherein we compared the full HIDE algorithm to two reduced versions of HIDE:

- **Ablation model 1:** in this model the gene-weight learning is performed directly on the sub-minor cell-types, without algorithmic additions. Respective estimates for major and minor cell types were derived by aggregating the predictions of the sub-minor cell types.
- **Ablation model 2:** in this model, we start with the base model of step (0) and then incorporate the hierarchical structure by training gene weights for each set of child cell types individually, corresponding to step (2) of the algorithm.

First, we compared the performance using the cell-type prediction at finest resolution, capturing in total 34 cell types. Results in terms of Pearson’s correlation were comparable, with values of 0.736±0.002 for HIDE, 0.739±0.002 for ablation model 1 and 0.725±0.002 for ablation model 2. Considering the corresponding NMAE values, however, the results were clearly in favor of HIDE, with an average NMAE of 0.086 ± 0.002 compared to 0.104 ± 0.003 for ablation model 1 and 0.258 ± 0.007 for ablation model 2 (see Table 3 and Table S14 and S17 for more detailed results). Thus, ablation model 2 has an approximately three-fold higher absolute error than HIDE, suggesting that steps (1) and (3) are essential for its predictions to also be quantitatively accurate. This may be driven by the fact that missing contributions in cell-type deconvolution inflate estimates of cellular proportions, as noted in (Nguyen *et al*., 2024; Sutton *et al*., 2022). In summary, at this resolution level, performance gains over ablation models in terms of Pearson’s correlation are minimal and mainly attributed to the gene-weight learning, whereas NMAE values are consistently lower for HIDE, suggesting better quantitative predictions.

**Table 3:**
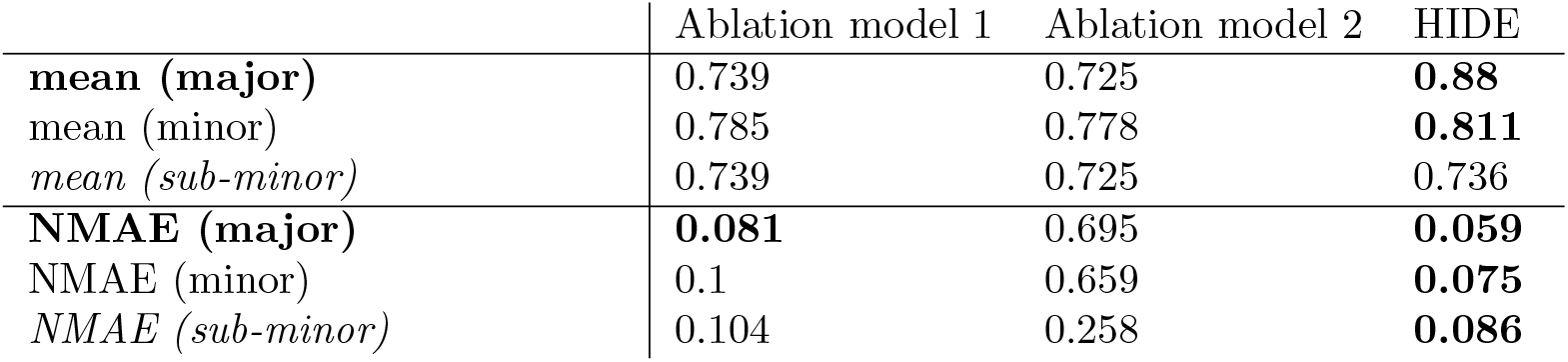
Performance comparison between HIDE and both Ablation models. Pearson’s correlations as well as NMAE values were generated by comparing predicted versus true cellular proportions for artificial test mixtures. Results correspond to the mean correlation obtained over 10 simulation runs. Correlations above 0.8 are highlighted in bold for Pearson’s correlation and below 0.1 for NMAE values.

Next, we aggregated the sub-minor cell type level predictions from HIDE and the two ablation models to retrieve corresponding predictions for minor and major cell types (Table 3). At the minor cell type level, results were again strongly in favor of HIDE in terms of NMAE, but also for Pearson’s correlation. HIDE achieved a Pearson’s correlation of 0.811 ± 0.002 compared to 0.785 ± 0.004 for ablation model 1 and 0.778 ± 0.002 for ablation model 2 (Table S15) and a mean NMAE value of 0.075 ± 0.003 compared to 0.1± 0.003 for ablation 1 and even 0.659 ± 0.026 for ablation 2 (Table S18). Results for the prediction of major cell types were in line with these findings with a Pearson’s correlation of 0.88 ± 0.003 for HIDE compared to 0.855 ± 0.004 for ablation model 1 and 0.833±0.004 for ablation model 2 (Table S16). Also here the NMAE values were lowest for HIDE with a mean NMAE value of 0.059 ± 0.002 compared to 0.081 ± 0.003 for ablation model 1 and 0.695 ± 0.028 for ablation model 2 (see Table S19).

These results illustrate that the combination of steps (1) to (3) are also essential to conserve estimates for larger cell populations. In other words, if cell-type deconvolution is performed for a high number of cell types, it might be essential to conserve estimates of cell groups, corresponding to minor and major cell types in our analysis.

### 3.3 Breast cancer subtype analysis using bulk transcriptomics data from TCGA

Finally, we compared HIDE with BayesPrism, CIBERSORTx and MuSiC in an application to bulk breast cancer expression profiles from TCGA. Models were established using breast cancer single-cell RNA-seq data obtained from DISCO (see Methods). The predicted cell type proportions were analyzed for their association with patient survival via Cox regression, employing the CoxPHFitter function from lifelines 0.30.0 (https://lifelines.readthedocs.io/en/latest/) with robust errors to account for possible violations of the model assumptions (Lin and Wei, 1989). Specifically, we performed survival analysis for the complete cohort as well as within individual breast cancer subtypes using overall survival as an end point.

#### 3.3.1 Cell type association with survival

To begin, we assessed the utility of each algorithm by determining the total number of corresponding cell type estimates significantly associated with survival via Cox regression analysis. The underlying rationale is that an association analysis is more likely to return significant findings when the covariates (in this case, cell type proportions) are more reliably predicted. Since the other competing algorithms do not provide an inherent hierarchical structure, we predicted the corresponding cellular proportions on the minor and major level via both the summing and the independent level approaches, as done previously. The number of significant cell types at different layers of resolution are summarized in Table S57 in the supplement. Corresponding detailed results including hazard ratios and *p*-values can be found in Tables S20-S56 for all four algorithms. At the major cell type level, HIDE predictions yielded nine significant cell types across all breast cancer subtypes; a result surpassed only by MuSiC, which identified ten significant cell types using the independent level approach. Meanwhile, BayesPrism yielded the fewest significant cell types with only four hits (summing approach). At the level of minor cell types, HIDE outperformed the other methods, yielding a total of 13 significant cell types, followed by MuSic using the independent level approach with 12 significant findings. At the finest resolution (sub-minor cell types), both HIDE and MuSic identified 27 significant cell types, followed by BayesPrism with 23 and CIBERSORTx with 22.

#### 3.3.2 Consistency of results

To assess the consistency of predictions across cell type levels, we evaluated cell types that were not subdivided into child categories (e.g., NK cells at all levels, endothelial cells at the major and minor level). We hypothesized that if a specific cell type is significantly associated with an outcome (e.g., shorter survival) at one level, it should exhibit a consistent association at all other levels.

To quantify this, we compared associations across major-minor, minor-sub-minor, and major-sub-minor levels for cell types retained in each pair of levels. An association in one level was considered consistent to another level if: (1) the cell type was significantly associated at both levels, and (2) the direction of the association (indicated by the coefficient of the hazard ratio) was the same at both levels. If one of these conditions was violated, we noted this association as inconsistent. We found for BayesPrism 3 inconsistent and 4 consistent, for CIBERSORTx 3 inconsistent and 5 consistent and for MuSiC 7 inconsistent and 7 consistent associations. Due to the definition of the summing approach it is impossible to find inconsistencies between different cell type levels. Such inconsistencies were also not observed for HIDE due to its top-down structure. Table S58 provides an overview over the findings.

Explicit findings are, for instance, that for BayesPrism, NK cells in triple-negative breast cancer patients were significant at both the major and minor levels, but not at the sub-minor level. Moreover, the sign of the coefficient changed from negative at the major cell level to positive at the minor cell level. A second example are proliferating T/NK cells in the Luminal B cancer subtype, which are hits for CIBERSORTx at the minor level, but not on the major or sub-minor level. Similarly, proliferating T/NK cells were found to be significant at the sub-minor level in the Luminal A and B cancer subtypes but not at the major level for CIBERSORTx.

To further investigate the consistency of the predictions, we identified “significant chains”, defined as instances where both a parent cell type and at least one of its child cell types were found to be significant predictors of patient survival in the Cox proportional hazards (PH) model. The number of significant chains within the hierarchical cell type structure was used to evaluate the performance of each deconvolution method. When a parent cell type as well as two of the corresponding child cell types show significant p-values in the Cox PH model, we counted 1 on the parent and 2 on the child level. The results are presented in Table S60 and S61 in the supplement for both the individual level and the summing approach, respectively. HIDE consistently demonstrated superior performance, exhibiting the highest number of significant chains at the minor and sub-minor-levels. While MuSiC identified a greater number of significant chains at the major level using the independent level approach, HIDE maintained the highest overall count of significant chains across all hierarchical levels. Interestingly, for BayesPrism and MuSiC, the individual level approach yielded a higher number of significant chains compared to the summing approach, despite the previously observed inconsistencies between these approaches. Compared to the summing approach, the independent level approach identified 3 and 4 more significant chains for BayesPrism and MuSiC, respectively. Conversely, CIBERSORTx identified 10 more significant chains with the summing approach.

#### 3.3.3 Biological Discussion of HIDE estimates

In total, 37 significant results (*p <* 0.05) were identified in the Cox regression analysis using the cellular proportions predicted by HIDE, including 5 in patients with the triple negative breast cancer subtype, 10 in HER2-positive breast cancer, 6 in luminal A, 8 in luminal B, and 8 in cases where breast cancer subtypes were not differentiated. Detailed results, including the Cox PH model coefficients and corresponding p-values, are provided in Tables S20 - S25.

Natural killer (NK) cells demonstrated a negative coefficient in the Cox PH model across all breast cancer subtypes, indicating a correlation between higher NK cell presence and improved overall survival (TNBC: *p* = 0.07, HER2: *p* = 0.037, LumA: *p* = 0.19, LumB: *p* = 0.02). NK cells are known to detect tumor-specific changes, including the loss of MHC class I, facilitating cancer cell elimination (Vivier *et al*., 2008). Consistent with this, (Liu *et al*., 2023) reported that triple-negative breast cancer patients with specific NK cell subtypes exhibited higher survival rates and better responses to therapy. Additionally, the Cox PH model associated CXCL13-exhausted CD8 positive T cells with improved overall survival (TNBC: *p* = 0.0036, HER2: *p* = 0.31, LumA: *p* = 0.036, LumB: *p* = 0.15). This aligns with findings by Lin et al., who observed that exhausted T cells expressing high levels of CXCL13 promote the formation of tertiary lymphoid structures (TLS) in breast cancer patients. TLS facilitate antigen presentation and lymphoid cell recruitment within tumors, contributing to enhanced survival outcomes (Lin *et al*., 2024). Furthermore, the impact of CXCL13-exhausted CD8 T cells on survival may also result from their interaction with immunotherapies. Zhang et al. demonstrated that these cells can enhance therapeutic responses (Zhang *et al*., 2021). In contrast, T-regulatory (Treg) cells were associated with poor prognosis in patients with the triple-negative breast cancer subtype (*p* = 0.049). These cells can suppress immune activity, thereby impairing the immune response against cancer cells (Adeegbe and Nishikawa, 2013; Elkord et al., 2010). The positive coefficient observed in the Cox PH model corroborates their immunosuppressive function and its association with reduced overall survival.

In addition, we investigated differences in the predicted tumor microenvironment of patients with only a primary tumor (TNM classification: *T >* 0, *N* = 0, *M* = 0) and patients with lymph node infiltration and metastasis (TNM classification: *T >* 0, *N >* 0, *M >* 0). Unknown degrees of lymph node infiltration (NX) and metastasis (MX) are considered as existing lymph node infiltration and metastasis. Since splitting the data first into breast cancer subtypes and then into the different TNM groups leads to small cohorts, we decided to omit the breast cancer subtypes and only examined the difference between the TNM groups. The p-values of the Cox PH regression analysis are shown in Table S26 in the supplement. In a mouse model, Gu et al. discovered that B cells can lead to lymph node infiltration and the production of pathogenic IgG antibodies, which is also consistent with our findings that the proportion of B cells is higher in patients with lymph node infiltration and metastasis than in patients with only a primary tumor (*p* = 0.0059) Gu et al. (2019). The Kruskal-Wallis test also revealed that CXCL 1*/*2*/*3 fibroblasts have lower expression levels in patients with only a primary tumor than in patients with lymph node invasion and metastasis (*p* = 0.0038). In mouse models, CXCL1 has been shown to be involved in various processes leading to aggressive invasive tumors. Also, suppression of TGF-*β* seems to lead to higher CXCL1 expression, which appears to be associated with metastases in breast cancer Bernard et al. (2018).

## 4 Conclusion and discussion

Cell-type deconvolution for bulk transcriptomics data has become a rapidly evolving research field with numerous methodological contributions in recent years (Cobos *et al*., 2023; Im and Kim, 2023). Meanwhile, emerging methods provide an increasingly fine-grained picture of cellular compositions, while also capturing cell populations which are highly underrepresented in bulk profiles. However, an inherent problem of reference based methods is that the number of investigated cell types substantially affects model results, as additional cell types add uncertainties to the underlying optimization problem. This is driven by the fact that new model parameters – the cellular proportions of additional cell types – have to be inferred from the data, which in turn affects estimates for the remaining cell types. Thus, new approaches are essential to increase the scope of cell-type deconvolution to the analysis of a large number of cell types, distinguishing molecularly related cell types and quantifying rare cell populations reliably. Importantly, these strategies should be designed such that results for larger cell populations remain unaffected, and not compromised as a consequence of the increased model complexity. HIDE can be considered as an important step in this direction. HIDE pays attention to the fact that cells can be structured as a hierarchy, resembling, for instance, cell lineages or phenotypic spectrums. This hierarchy is incorporated as a criterion to stabilize estimates. In other words, HIDE constrains its model parameters – the cellular proportions – to be consistent with a user-provided hierarchy. As a consequence, smaller cell populations, such as pericytes and smooth muscle cells, sum up to their corresponding parent cell type (perivascular cells). Thus, in case that one or more sub-populations cannot be inferred accurately, the upper layers remain unaffected, limiting the negative effect of increased model complexity. Moreover, HIDE makes rigorous use of the available gene space. We have repeatedly shown that optimized gene weights are highly beneficial for cell-type deconvolution (Görtler *et al*., 2020; Görtler et al., 2024). HIDE adapts this concept to accommodate optimized deconvolution at each layer of the cell hierarchy.

HIDE has limitations, which need to be addressed in future work. First, cell-type deconvolution can be affected by unseen contributions in the training data. To address this issue, background contributions could be directly incorporated into the optimization problem (Görtler *et al*., 2024; Mensching-Buhr *et al*., 2024). Moreover, the scope of cell-type deconvolution increased substantially in recent years, with methods such as BayesPrism, TissueResolver and ADTD (Chu *et al*., 2022; Simeth *et al*., 2024; Görtler *et al*., 2024) now able to facilitate estimates of cell-type specific gene regulation. Such capabilities could also be valuable extensions to HIDE.

In summary, HIDE provides a state-of-the-art framework for cell-type deconvolution which makes use of the hierarchical structure of cellular populations. It systematically enhances the scope of cell-type deconvolution to substantially more cell types, being able to resolve also underrepresented and molecularly related cell types reliably.

