## Supplement for "HIDE: Hierarchical cell-type Deconvolution"

### Supplementary Material: HIDE

#### 1 Informations corresponding to the single cell RNA-Seq dataset

| | single cells in training set | single cells in test set | single cells in BayesPrism | percentage |
|---|---|---|---|---|
| Arterial EC | 340 | 341 | 170 | 0.3 |
| Breast basal cell | 1741 | 1741 | 870 | 1.8 |
| Breast cancer specific luminal cell | 26321 | 26321 | 13160 | 26.7 |
| Breast cancer specific proliferation luminal cell | 3307 | 3308 | 1653 | 3.4 |
| Capillary EC | 1014 | 1014 | 507 | 1 |
| CD4 T | 7716 | 7717 | 3858 | 7.8 |
| cDC2 | 1166 | 1166 | 583 | 1.2 |
| CFD fibroblast | 1582 | 1583 | 791 | 1.6 |
| CXCL1/2/3 fibroblast | 2847 | 2848 | 1423 | 2.9 |
| CXCL13 exhausted CD8 T | 1146 | 1146 | 573 | 1.2 |
| GZMH CD8 T | 3893 | 3894 | 1946 | 4 |
| GZMK CD8 T | 4952 | 4952 | 2476 | 5 |
| IgA plasma | 569 | 569 | 284 | 0.6 |
| IgG plasma cell | 2241 | 2241 | 1120 | 2.3 |
| INF responsed T | 97 | 97 | 48 | 0.1 |
| Luminal progenitor | 2646 | 2646 | 1323 | 2.7 |
| Lymphatic EC | 162 | 162 | 81 | 0.2 |
| Macrophage | 7427 | 7428 | 3713 | 7.5 |
| Mast cell | 444 | 444 | 222 | 0.5 |
| Monocyte | 1120 | 1120 | 560 | 1.1 |
| mregDC | 134 | 134 | 67 | 0.1 |
| NK cell | 1594 | 1595 | 797 | 1.6 |
| Other B cells | 2514 | 2515 | 1257 | 2.6 |
| Other fibroblasts | 3576 | 3577 | 1788 | 3.6 |
| pDC | 325 | 325 | 162 | 0.3 |
| Pericyte | 472 | 472 | 236 | 0.5 |
| Proliferation macrophage | 507 | 507 | 253 | 0.5 |
| Proliferation T/NK | 1053 | 1053 | 526 | 1.1 |
| Smooth muscle cell | 1487 | 1487 | 743 | 1.5 |
| Tfh | 2070 | 2070 | 1035 | 2.1 |
| TGM2 luminal cell | 723 | 723 | 361 | 0.7 |
| TNBC-specific epithelial cell | 8496 | 8496 | 4248 | 8.6 |
| Treg | 3495 | 3496 | 1747 | 3.5 |
| Venous EC | 1276 | 1277 | 638 | 1.3 |
| sum | 98453 | 98465 | 49219 | 100 |

Table S1: **Distribution of the single cells used for benchmarking.** Distribution of single cells used to create the artificial training and test set and the number of single cells used for training with BayesPrism. The percentage remains the same for training, test and BayesPrism set.

| | BayesPrism | CIBERSORTx | MuSiC | HIDE |
|---|---|---|---|---|
| Arterial EC | 0.445 ± 0.042 | 0.321 ± 0.041 | 0.345 ± 0.041 | 0.643 ± 0.033 |
| Breast basal cell | 0.561 ± 0.026 | 0.663 ± 0.022 | 0.433 ± 0.012 | **0.878 ± 0.005** |
| Breast cancer specific luminal cell | 0.531 ± 0.025 | 0.65 ± 0.025 | 0.656 ± 0.025 | **0.907 ± 0.006** |
| Breast cancer specific proliferation luminal cell | 0.311 ± 0.026 | 0.307 ± 0.038 | 0.348 ± 0.031 | 0.625 ± 0.02 |
| Capillary EC | 0.487 ± 0.03 | 0.459 ± 0.018 | 0.275 ± 0.037 | 0.719 ± 0.022 |
| CD4 T | 0.612 ± 0.025 | 0.475 ± 0.037 | 0.465 ± 0.023 | 0.769 ± 0.011 |
| cDC2 | 0.543 ± 0.017 | 0.4 ± 0.024 | 0.48 ± 0.022 | 0.689 ± 0.023 |
| CFD fibroblast | 0.565 ± 0.031 | 0.666 ± 0.016 | 0.501 ± 0.031 | **0.812 ± 0.01** |
| CXCL1/2/3 fibroblast | 0.696 ± 0.02 | 0.788 ± 0.012 | 0.536 ± 0.03 | **0.901 ± 0.008** |
| CXCL13 exhausted CD8 T | 0.396 ± 0.026 | 0.255 ± 0.026 | 0.296 ± 0.031 | 0.556 ± 0.015 |
| GZMH CD8 T | 0.424 ± 0.031 | 0.277 ± 0.028 | 0.177 ± 0.026 | 0.612 ± 0.022 |
| GZMK CD8 T | 0.626 ± 0.024 | 0.555 ± 0.029 | 0.366 ± 0.046 | 0.78 ± 0.015 |
| IgA plasma | 0.791 ± 0.01 | 0.75 ± 0.018 | 0.687 ± 0.018 | **0.877 ± 0.007** |
| IgG plasma cell | 0.757 ± 0.019 | 0.787 ± 0.011 | 0.775 ± 0.014 | **0.936 ± 0.004** |
| INF responsed T | 0.234 ± 0.048 | 0.206 ± 0.029 | 0.142 ± 0.03 | 0.176 ± 0.049 |
| Luminal progenitor | 0.565 ± 0.017 | 0.5 ± 0.021 | 0.484 ± 0.024 | **0.838 ± 0.005** |
| Lymphatic EC | 0.636 ± 0.033 | 0.488 ± 0.029 | 0.227 ± 0.03 | 0.763 ± 0.017 |
| Macrophage | 0.509 ± 0.027 | 0.586 ± 0.02 | 0.505 ± 0.024 | **0.898 ± 0.005** |
| Mast cell | 0.599 ± 0.028 | 0.584 ± 0.028 | 0.523 ± 0.036 | 0.745 ± 0.019 |
| Monocyte | 0.266 ± 0.03 | 0.209 ± 0.04 | 0.263 ± 0.037 | 0.383 ± 0.018 |
| mregDC | 0.395 ± 0.045 | 0.343 ± 0.035 | 0.368 ± 0.044 | 0.651 ± 0.028 |
| NK cell | 0.524 ± 0.02 | 0.4 ± 0.024 | 0.416 ± 0.021 | 0.757 ± 0.01 |
| Other B cells | 0.514 ± 0.03 | 0.492 ± 0.034 | 0.429 ± 0.033 | **0.84 ± 0.009** |
| Other fibroblasts | 0.706 ± 0.019 | **0.843 ± 0.014** | 0.64 ± 0.021 | **0.909 ± 0.007** |
| pDC | 0.624 ± 0.035 | 0.571 ± 0.025 | 0.447 ± 0.027 | **0.831 ± 0.017** |
| Pericyte | 0.327 ± 0.037 | 0.4 ± 0.028 | 0.338 ± 0.04 | 0.626 ± 0.031 |
| Proliferation macrophage | 0.219 ± 0.028 | 0.16 ± 0.036 | 0.206 ± 0.043 | 0.38 ± 0.035 |
| Proliferation T/NK | 0.271 ± 0.027 | 0.267 ± 0.034 | 0.189 ± 0.052 | 0.574 ± 0.027 |
| Smooth muscle cell | 0.588 ± 0.017 | 0.69 ± 0.017 | 0.234 ± 0.033 | **0.865 ± 0.01** |
| Tfh | 0.554 ± 0.016 | 0.574 ± 0.021 | 0.515 ± 0.015 | 0.736 ± 0.01 |
| TGM2 luminal cell | 0.342 ± 0.034 | 0.369 ± 0.044 | 0.149 ± 0.045 | 0.735 ± 0.017 |
| TNBC-specific epithelial cell | 0.769 ± 0.011 | 0.897 ± 0.006 | 0.517 ± 0.025 | **0.945 ± 0.005** |
| Treg | 0.622 ± 0.025 | 0.566 ± 0.033 | 0.453 ± 0.014 | **0.818 ± 0.011** |
| Venous EC | 0.597 ± 0.024 | 0.568 ± 0.03 | 0.544 ± 0.023 | **0.839 ± 0.008** |
| mean | 0.518 ± 0.003 | 0.502 ± 0.006 | 0.41 ± 0.004 | 0.736 ± 0.002 |

Table S2: **Observed Pearson's correlations on sub-minor level,** by comparing the estimated cell proportions with the true cell proportions predicted in the benchmarking study for artificial test mixtures. The error bars correspond to ±1 SD obtained over 10 simulation runs. Correlations above 0.8 are highlighted in bold.

| | BayesPrism | CIBERSORTx | MuSiC | HIDE |
|---|---|---|---|---|
| B cell | 0.653 ± 0.028 | 0.646 ± 0.027 | 0.629 ± 0.028 | **0.931 ± 0.005** |
| CD4 T cell | 0.696 ± 0.019 | 0.622 ± 0.027 | 0.604 ± 0.025 | **0.884 ± 0.006** |
| CD8 T cell | 0.614 ± 0.018 | 0.494 ± 0.02 | 0.453 ± 0.026 | **0.838 ± 0.006** |
| Dendritic cell | 0.534 ± 0.021 | 0.445 ± 0.031 | 0.476 ± 0.025 | 0.754 ± 0.02 |
| Endothelial cell | 0.724 ± 0.009 | 0.734 ± 0.007 | 0.654 ± 0.018 | **0.946 ± 0.003** |
| Epithelial cell | 0.685 ± 0.019 | **0.865 ± 0.01** | 0.711 ± 0.015 | **0.962 ± 0.003** |
| Fibroblast | 0.701 ± 0.02 | **0.866 ± 0.008** | 0.664 ± 0.022 | **0.953 ± 0.003** |
| Granulocyte | 0.599 ± 0.028 | 0.584 ± 0.028 | 0.523 ± 0.036 | 0.745 ± 0.019 |
| Macrophage | 0.552 ± 0.015 | 0.628 ± 0.019 | 0.533 ± 0.02 | **0.909 ± 0.006** |
| Monocyte | 0.266 ± 0.03 | 0.209 ± 0.04 | 0.263 ± 0.037 | 0.383 ± 0.018 |
| NK cell | 0.524 ± 0.02 | 0.4 ± 0.024 | 0.416 ± 0.021 | 0.757 ± 0.01 |
| Perivascular cell | 0.557 ± 0.022 | 0.708 ± 0.023 | 0.423 ± 0.022 | **0.905 ± 0.006** |
| Proliferation T/NK | 0.271 ± 0.027 | 0.267 ± 0.034 | 0.189 ± 0.052 | 0.574 ± 0.027 |
| mean | 0.567 ± 0.005 | 0.575 ± 0.006 | 0.503 ± 0.006 | **0.811 ± 0.002** |

Table S3: **Observed Pearson's correlations on minor level predicted via summing approach.** Pearson's correlations are calculated between true and predited cellular composition. The error bars correspond to ±1 SD obtained over 10 simulation runs. Correlations above 0.8 are highlighted in bold.

| | BayesPrism | CIBERSORTx | MuSiC | HIDE |
|---|---|---|---|---|
| B cell | 0.653 ± 0.028 | 0.646 ± 0.027 | 0.629 ± 0.028 | **0.931 ± 0.005** |
| Endothelial cell | 0.724 ± 0.009 | 0.734 ± 0.007 | 0.654 ± 0.018 | **0.946 ± 0.003** |
| Epithelial cell | 0.685 ± 0.019 | **0.865 ± 0.01** | 0.711 ± 0.015 | **0.962 ± 0.003** |
| Fibroblast | 0.701 ± 0.02 | **0.866 ± 0.008** | 0.664 ± 0.022 | **0.953 ± 0.003** |
| Myeloid cell | 0.666 ± 0.022 | 0.757 ± 0.014 | 0.655 ± 0.021 | **0.965 ± 0.002** |
| NK cell | 0.524 ± 0.02 | 0.4 ± 0.024 | 0.416 ± 0.021 | 0.757 ± 0.01 |
| Perivascular cell | 0.557 ± 0.022 | 0.708 ± 0.023 | 0.423 ± 0.022 | **0.905 ± 0.006** |
| Proliferation T/NK | 0.271 ± 0.027 | 0.267 ± 0.034 | 0.189 ± 0.052 | 0.574 ± 0.027 |
| T cell | 0.707 ± 0.02 | 0.701 ± 0.023 | 0.706 ± 0.016 | **0.924 ± 0.003** |
| mean | 0.61 ± 0.007 | 0.661 ± 0.007 | 0.561 ± 0.006 | **0.88 ± 0.003** |

Table S4: **Observed Pearson's correlations on major level predicted via summing approach.** Pearson's correlations are calculated between true and predited cellular composition. The error bars correspond to ±1 SD obtained over 10 simulation runs. Correlations above 0.8 are highlighted in bold.

| | BayesPrism | CIBERSORTx | MuSiC | HIDE |
|---|---|---|---|---|
| B cell | 0.507 ± 0.029 | 0.6 ± 0.028 | 0.63 ± 0.029 | **0.931 ± 0.005** |
| CD4 T cell | 0.659 ± 0.024 | 0.572 ± 0.022 | 0.51 ± 0.037 | **0.884 ± 0.006** |
| CD8 T cell | 0.564 ± 0.017 | 0.553 ± 0.028 | 0.47 ± 0.028 | **0.838 ± 0.006** |
| Dendritic cell | 0.508 ± 0.028 | 0.343 ± 0.039 | 0.434 ± 0.024 | 0.754 ± 0.02 |
| Endothelial cell | 0.66 ± 0.016 | 0.605 ± 0.019 | 0.588 ± 0.02 | **0.946 ± 0.003** |
| Epithelial cell | 0.657 ± 0.019 | **0.865 ± 0.007** | 0.689 ± 0.015 | **0.962 ± 0.003** |
| Fibroblast | 0.663 ± 0.022 | **0.819 ± 0.011** | 0.598 ± 0.023 | **0.953 ± 0.003** |
| Granulocyte | 0.598 ± 0.026 | 0.28 ± 0.037 | 0.504 ± 0.035 | 0.745 ± 0.019 |
| Macrophage | 0.546 ± 0.015 | 0.612 ± 0.02 | 0.542 ± 0.015 | **0.909 ± 0.006** |
| Monocyte | 0.255 ± 0.025 | 0.187 ± 0.027 | 0.28 ± 0.034 | 0.383 ± 0.018 |
| NK cell | 0.195 ± 0.04 | 0.34 ± 0.024 | 0.392 ± 0.027 | 0.757 ± 0.01 |
| Perivascular cell | 0.601 ± 0.016 | 0.642 ± 0.022 | 0.424 ± 0.034 | **0.905 ± 0.006** |
| Proliferation T/NK | 0.224 ± 0.039 | 0.307 ± 0.041 | 0.16 ± 0.051 | 0.574 ± 0.027 |
| mean | 0.51 ± 0.006 | 0.517 ± 0.007 | 0.479 ± 0.006 | **0.811 ± 0.002** |

Table S5: **Observed Pearson's correlations on minor level predicted via individual level approach.** Pearson's correlations are calculated between true and predited cellular composition. The error bars correspond to ±1 SD obtained over 10 simulation runs. Correlations above 0.8 are highlighted in bold.

| | BayesPrism | CIBERSORTx | MuSiC | HIDE |
|---|---|---|---|---|
| B cell | 0.512 ± 0.029 | 0.598 ± 0.028 | 0.636 ± 0.029 | **0.931 ± 0.005** |
| Endothelial cell | 0.659 ± 0.015 | 0.603 ± 0.019 | 0.595 ± 0.018 | **0.946 ± 0.003** |
| Epithelial cell | 0.662 ± 0.019 | **0.878 ± 0.007** | 0.686 ± 0.015 | **0.962 ± 0.003** |
| Fibroblast | 0.665 ± 0.022 | **0.822 ± 0.011** | 0.599 ± 0.023 | **0.953 ± 0.003** |
| Myeloid cell | 0.66 ± 0.023 | **0.863 ± 0.008** | 0.657 ± 0.019 | **0.965 ± 0.002** |
| NK cell | 0.399 ± 0.021 | 0.273 ± 0.022 | 0.298 ± 0.019 | 0.757 ± 0.01 |
| Perivascular cell | 0.599 ± 0.016 | 0.603 ± 0.023 | 0.423 ± 0.031 | **0.905 ± 0.006** |
| Proliferation T/NK | 0.231 ± 0.042 | 0.299 ± 0.045 | 0.166 ± 0.053 | 0.574 ± 0.027 |
| T cell | 0.653 ± 0.021 | 0.668 ± 0.016 | 0.539 ± 0.028 | **0.924 ± 0.003** |
| mean | 0.56 ± 0.008 | 0.623 ± 0.006 | 0.511 ± 0.01 | **0.88 ± 0.003** |

Table S6: **Observed Pearson's correlations on major level predicted via individual level approach.** Pearson's correlations are calculated between true and predited cellular composition. The error bars correspond to ±1 SD obtained over 10 simulation runs. Correlations above 0.8 are highlighted in bold.

| | BayesPrism | CIBERSORTx | MuSiC | HIDE |
|---|---|---|---|---|
| B cell | 0.594 ± 0.028 | 0.575 ± 0.021 | 0.639 ± 0.024 | **1** |
| CD4 T cell | **0.88 ± 0.007** | **0.805 ± 0.01** | **0.863 ± 0.01** | **1** |
| CD8 T cell | 0.79 ± 0.019 | 0.663 ± 0.015 | 0.794 ± 0.011 | **1** |
| Dendritic cell | **0.836 ± 0.016** | 0.588 ± 0.027 | **0.911 ± 0.009** | **1** |
| Endothelial cell | **0.87 ± 0.006** | 0.63 ± 0.017 | **0.928 ± 0.006** | **1** |
| Epithelial cell | **0.955 ± 0.002** | **0.854 ± 0.01** | **0.968 ± 0.002** | **1** |
| Fibroblast | **0.963 ± 0.003** | **0.877 ± 0.008** | **0.94 ± 0.003** | **1** |
| Granulocyte | **0.995 ± 0.001** | 0.678 ± 0.021 | **0.908 ± 0.013** | **1** |
| Macrophage | **0.978 ± 0.002** | 0.87 ± 0.009 | **0.982 ± 0.001** | **1** |
| Monocyte | **0.868 ± 0.01** | 0.58 ± 0.027 | **0.94 ± 0.004** | **1** |
| NK cell | 0.355 ± 0.05 | 0.558 ± 0.025 | 0.78 ± 0.015 | **1** |
| Perivascular cell | 0.795 ± 0.018 | 0.765 ± 0.02 | 0.794 ± 0.028 | **1** |
| Proliferation T/NK | 0.426 ± 0.032 | 0.308 ± 0.03 | 0.598 ± 0.019 | **1** |
| mean | 0.793 ± 0.008 | 0.673 ± 0.007 | **0.85 ± 0.005** | 1 |

Table S7: **Observed Pearson's correlations on the minor cellular level resulting form comparing summing and individual level approach.** The error bars correspond to ±1 SD obtained over 10 simulation runs. Correlations above 0.8 are highlighted in bold.

| | BayesPrism | CIBERSORTx | MuSiC | HIDE |
|---|---|---|---|---|
| B cell | 0.603 ± 0.028 | 0.575 ± 0.02 | 0.647 ± 0.023 | **1** |
| Endothelial cell | 0.868 ± 0.006 | 0.626 ± 0.019 | **0.92 ± 0.006** | **1** |
| Epithelial cell | **0.96 ± 0.002** | 0.853 ± 0.01 | **0.969 ± 0.002** | **1** |
| Fibroblast | **0.965 ± 0.003** | 0.873 ± 0.008 | **0.942 ± 0.004** | **1** |
| Myeloid cell | **0.973 ± 0.001** | 0.803 ± 0.008 | **0.959 ± 0.002** | **1** |
| NK cell | 0.638 ± 0.028 | 0.343 ± 0.034 | 0.599 ± 0.019 | **1** |
| Perivascular cell | 0.791 ± 0.018 | 0.737 ± 0.022 | 0.795 ± 0.026 | **1** |
| Proliferation T/NK | 0.45 ± 0.031 | 0.261 ± 0.033 | 0.59 ± 0.019 | **1** |
| T cell | 0.889 ± 0.008 | 0.715 ± 0.015 | 0.769 ± 0.014 | **1** |
| mean | 0.793 ± 0.007 | 0.643 ± 0.009 | 0.799 ± 0.007 | **1** |

Table S8: **Observed Pearson's correlations on major level resulting form comparing summing and individual level approach.** The error bars correspond to ±1 SD obtained over 10 simulation runs. Correlations above 0.8 are highlighted in bold.

##### 1.1 Nmae results

| | BayesPrism | CIBERSORTx | MuSiC | HIDE |
|---|---|---|---|---|
| Arterial EC | 0.129 ± 0.017 | 0.209 ± 0.03 | 0.126 ± 0.017 | **0.089 ± 0.012** |
| Breast basal cell | 0.145 ± 0.016 | 0.132 ± 0.015 | 0.331 ± 0.051 | **0.07 ± 0.01** |
| Breast cancer specific luminal cell | 0.208 ± 0.017 | 0.159 ± 0.014 | 0.295 ± 0.024 | **0.053 ± 0.004** |
| Breast cancer specific proliferation luminal cell | 0.38 ± 0.045 | 0.276 ± 0.034 | 0.646 ± 0.064 | 0.107 ± 0.01 |
| Capillary EC | 0.134 ± 0.015 | 0.184 ± 0.024 | 0.165 ± 0.019 | 0.103 ± 0.012 |
| CD4 T | 0.21 ± 0.02 | 0.209 ± 0.021 | 0.506 ± 0.054 | **0.084 ± 0.008** |
| cDC2 | 0.245 ± 0.021 | 0.208 ± 0.017 | 0.242 ± 0.021 | 0.118 ± 0.012 |
| CFD fibroblast | 0.147 ± 0.023 | 0.124 ± 0.016 | 0.172 ± 0.026 | **0.084 ± 0.011** |
| CXCL1/2/3 fibroblast | 0.131 ± 0.017 | 0.1 ± 0.011 | 0.141 ± 0.015 | **0.058 ± 0.007** |
| CXCL13 exhausted CD8 T | 0.157 ± 0.019 | 0.264 ± 0.035 | 0.627 ± 0.075 | 0.131 ± 0.014 |
| GZMH CD8 T | 0.23 ± 0.023 | 0.343 ± 0.036 | 0.313 ± 0.033 | 0.111 ± 0.01 |
| GZMK CD8 T | 0.185 ± 0.021 | 0.163 ± 0.018 | 0.256 ± 0.025 | **0.086 ± 0.01** |
| IgA plasma | 0.123 ± 0.016 | **0.078 ± 0.011** | **0.094 ± 0.014** | **0.06 ± 0.009** |
| IgG plasma cell | 0.102 ± 0.012 | **0.098 ± 0.012** | 0.121 ± 0.015 | **0.052 ± 0.006** |
| INF responsed T | 0.383 ± 0.06 | 0.262 ± 0.041 | 0.836 ± 0.13 | 0.101 ± 0.014 |
| Luminal progenitor | 0.186 ± 0.017 | 0.181 ± 0.018 | 0.246 ± 0.027 | **0.076 ± 0.008** |
| Lymphatic EC | **0.073 ± 0.013** | 0.125 ± 0.022 | 0.127 ± 0.025 | **0.059 ± 0.011** |
| Macrophage | 0.221 ± 0.011 | 0.165 ± 0.011 | 0.236 ± 0.012 | **0.059 ± 0.004** |
| Mast cell | **0.089 ± 0.01** | 0.123 ± 0.017 | 0.106 ± 0.012 | **0.094 ± 0.013** |
| Monocyte | 0.479 ± 0.066 | 0.399 ± 0.045 | 0.541 ± 0.069 | **0.143 ± 0.018** |
| mregDC | 0.145 ± 0.031 | 0.176 ± 0.033 | **0.07 ± 0.014** | **0.058 ± 0.012** |
| NK cell | 0.147 ± 0.016 | 0.214 ± 0.026 | 0.271 ± 0.03 | **0.097 ± 0.011** |
| Other B cells | 0.201 ± 0.025 | 0.192 ± 0.028 | 0.279 ± 0.036 | **0.082 ± 0.011** |
| Other fibroblasts | 0.124 ± 0.009 | **0.083 ± 0.007** | 0.191 ± 0.014 | **0.057 ± 0.004** |
| pDC | **0.088 ± 0.013** | 0.137 ± 0.018 | 0.17 ± 0.023 | **0.066 ± 0.01** |
| Pericyte | 0.163 ± 0.027 | 0.169 ± 0.029 | 0.137 ± 0.025 | **0.098 ± 0.018** |
| Proliferation macrophage | 0.513 ± 0.047 | 0.352 ± 0.027 | 0.137 ± 0.015 | 0.139 ± 0.013 |
| Proliferation T/NK | 0.202 ± 0.022 | 0.302 ± 0.035 | 0.21 ± 0.022 | 0.13 ± 0.014 |
| Smooth muscle cell | 0.149 ± 0.018 | 0.119 ± 0.014 | 0.223 ± 0.027 | **0.073 ± 0.009** |
| Tfh | 0.159 ± 0.022 | 0.161 ± 0.02 | 0.248 ± 0.036 | **0.099 ± 0.012** |
| TGM2 luminal cell | 0.225 ± 0.032 | 0.233 ± 0.03 | 0.178 ± 0.023 | **0.093 ± 0.012** |
| TNBC-specific epithelial cell | 0.312 ± 0.027 | **0.072 ± 0.006** | 0.343 ± 0.029 | **0.04 ± 0.004** |
| Treg | 0.163 ± 0.013 | 0.163 ± 0.015 | 0.23 ± 0.016 | **0.079 ± 0.006** |
| Venous EC | 0.127 ± 0.02 | 0.145 ± 0.024 | 0.143 ± 0.02 | **0.079 ± 0.012** |
| mean | 0.196 ± 0.005 | 0.186 ± 0.004 | 0.263 ± 0.007 | **0.086 ± 0.002** |

Table S9: **NMAE error on sub cellular level.** Received by comparing the estimated cell proportions with the true cell proportions on the artificial test mixtures. The error bars correspond to ±1 SD obtained over 10 simulation runs. Values below 0.1 are highlighted in bold.

| | BayesPrism | CIBERSORTx | MuSiC | HIDE |
|---|---|---|---|---|
| B cell | 0.146 ± 0.013 | 0.131 ± 0.012 | 0.183 ± 0.017 | **0.047 ± 0.004** |
| CD4 T cell | 0.209 ± 0.015 | 0.133 ± 0.01 | 0.38 ± 0.024 | **0.058 ± 0.004** |
| CD8 T cell | 0.265 ± 0.019 | 0.179 ± 0.011 | 0.205 ± 0.014 | **0.069 ± 0.005** |
| Dendritic cell | 0.226 ± 0.031 | 0.207 ± 0.032 | 0.231 ± 0.029 | **0.099 ± 0.015** |
| Endothelial cell | 0.113 ± 0.011 | 0.132 ± 0.012 | 0.134 ± 0.013 | **0.046 ± 0.005** |
| Epithelial cell | 0.297 ± 0.026 | 0.122 ± 0.011 | 0.176 ± 0.017 | **0.035 ± 0.003** |
| Fibroblast | 0.132 ± 0.011 | 0.087 ± 0.007 | 0.167 ± 0.014 | **0.038 ± 0.003** |
| Granulocyte | **0.089 ± 0.01** | 0.123 ± 0.017 | 0.106 ± 0.012 | **0.094 ± 0.013** |
| Macrophage | 0.219 ± 0.014 | 0.141 ± 0.009 | 0.237 ± 0.014 | **0.054 ± 0.004** |
| Monocyte | 0.479 ± 0.066 | 0.399 ± 0.045 | 0.541 ± 0.069 | 0.143 ± 0.018 |
| NK cell | 0.147 ± 0.016 | 0.214 ± 0.026 | 0.271 ± 0.03 | **0.097 ± 0.011** |
| Perivascular cell | 0.14 ± 0.015 | 0.114 ± 0.009 | 0.2 ± 0.022 | **0.061 ± 0.007** |
| Proliferation T/NK | 0.202 ± 0.022 | 0.302 ± 0.035 | 0.21 ± 0.022 | 0.13 ± 0.014 |
| mean | 0.205 ± 0.008 | 0.176 ± 0.006 | 0.234 ± 0.007 | **0.075 ± 0.003** |

Table S10: **NMAE values on minor cellular level predicted via summing approach.** Received by comparing the estimated cell proportions with the true cell proportions on the artificial test mixtures. The error bars correspond to ±1 SD obtained over 10 simulation runs. Values below 0.1 are highlighted in bold.

| | BayesPrism | CIBERSORTx | MuSiC | HIDE |
|---|---|---|---|---|
| B cell | 0.146 ± 0.013 | 0.131 ± 0.012 | 0.183 ± 0.017 | **0.047 ± 0.004** |
| Endothelial cell | 0.113 ± 0.011 | 0.132 ± 0.012 | 0.134 ± 0.013 | **0.046 ± 0.005** |
| Epithelial cell | 0.297 ± 0.026 | 0.122 ± 0.011 | 0.176 ± 0.017 | **0.035 ± 0.003** |
| Fibroblast | 0.132 ± 0.011 | **0.087 ± 0.007** | 0.167 ± 0.014 | **0.038 ± 0.003** |
| Myeloid cell | 0.235 ± 0.021 | 0.128 ± 0.013 | 0.164 ± 0.015 | **0.033 ± 0.003** |
| NK cell | 0.147 ± 0.016 | 0.214 ± 0.026 | 0.271 ± 0.03 | **0.097 ± 0.011** |
| Perivascular cell | 0.14 ± 0.015 | 0.114 ± 0.009 | 0.2 ± 0.022 | **0.061 ± 0.007** |
| Proliferation T/NK | 0.202 ± 0.022 | 0.302 ± 0.035 | 0.21 ± 0.022 | 0.13 ± 0.014 |
| T cell | 0.355 ± 0.024 | 0.111 ± 0.008 | 0.265 ± 0.017 | **0.047 ± 0.003** |
| mean | 0.196 ± 0.006 | 0.149 ± 0.005 | 0.197 ± 0.006 | **0.059 ± 0.002** |

Table S11: **NMAE values on major cellular level predicted via summing approach.** Received by comparing the estimated cell proportions with the true cell proportions on the artificial test mixtures. The error bars correspond to ±1 SD obtained over 10 simulation runs. Values below 0.1 are highlighted in bold.

| | BayesPrism | CIBERSORTx | MuSiC | HIDE |
|---|---|---|---|---|
| B cell | 0.218 ± 0.021 | 0.168 ± 0.016 | 0.214 ± 0.02 | **0.047 ± 0.004** |
| CD4 T cell | 0.27 ± 0.018 | 0.171 ± 0.012 | 0.293 ± 0.022 | **0.058 ± 0.004** |
| CD8 T cell | 0.287 ± 0.02 | 0.175 ± 0.015 | 0.283 ± 0.018 | **0.069 ± 0.005** |
| Dendritic cell | 0.213 ± 0.029 | 0.273 ± 0.043 | 0.224 ± 0.03 | **0.099 ± 0.015** |
| Endothelial cell | 0.132 ± 0.015 | 0.155 ± 0.017 | 0.139 ± 0.015 | **0.046 ± 0.005** |
| Epithelial cell | 0.319 ± 0.028 | **0.089 ± 0.009** | 0.231 ± 0.021 | **0.035 ± 0.003** |
| Fibroblast | 0.142 ± 0.012 | **0.093 ± 0.007** | 0.201 ± 0.017 | **0.038 ± 0.003** |
| Granulocyte | **0.089 ± 0.01** | 0.311 ± 0.047 | 0.126 ± 0.013 | **0.094 ± 0.013** |
| Macrophage | 0.208 ± 0.013 | 0.161 ± 0.01 | 0.219 ± 0.014 | **0.054 ± 0.004** |
| Monocyte | 0.535 ± 0.073 | 0.472 ± 0.063 | 0.566 ± 0.07 | 0.143 ± 0.018 |
| NK cell | 0.21 ± 0.027 | 0.281 ± 0.035 | 0.379 ± 0.042 | **0.097 ± 0.011** |
| Perivascular cell | 0.14 ± 0.016 | 0.146 ± 0.014 | 0.213 ± 0.024 | **0.061 ± 0.007** |
| Proliferation T/NK | 0.39 ± 0.046 | 0.29 ± 0.033 | 0.368 ± 0.045 | 0.13 ± 0.014 |
| mean | 0.242 ± 0.009 | 0.214 ± 0.009 | 0.266 ± 0.009 | **0.075 ± 0.003** |

Table S12: **NMAE values on minor cellular level predicted via independent level approach.** Received by comparing the estimated cell proportions with the true cell proportions on the artificial test mixtures. The error bars correspond to ±1 SD obtained over 10 simulation runs. Values below 0.1 are highlighted in bold.

| | BayesPrism | CIBERSORTx | MuSiC | HIDE |
|---|---|---|---|---|
| B cell | 0.208 ± 0.02 | 0.173 ± 0.016 | 0.217 ± 0.021 | **0.047 ± 0.004** |
| Endothelial cell | 0.131 ± 0.015 | 0.159 ± 0.018 | 0.137 ± 0.014 | **0.046 ± 0.005** |
| Epithelial cell | 0.273 ± 0.024 | **0.071 ± 0.007** | 0.245 ± 0.022 | **0.035 ± 0.003** |
| Fibroblast | 0.145 ± 0.013 | **0.091 ± 0.007** | 0.201 ± 0.017 | **0.038 ± 0.003** |
| Myeloid cell | 0.242 ± 0.021 | **0.075 ± 0.005** | 0.165 ± 0.014 | **0.033 ± 0.003** |
| NK cell | 0.164 ± 0.018 | 0.337 ± 0.04 | 0.425 ± 0.046 | **0.097 ± 0.011** |
| Perivascular cell | 0.141 ± 0.016 | 0.165 ± 0.015 | 0.214 ± 0.024 | **0.061 ± 0.007** |
| Proliferation T/NK | 0.382 ± 0.043 | 0.305 ± 0.035 | 0.382 ± 0.047 | 0.13 ± 0.014 |
| T cell | 0.386 ± 0.024 | 0.136 ± 0.01 | 0.33 ± 0.021 | **0.047 ± 0.003** |
| mean | 0.23 ± 0.007 | 0.168 ± 0.007 | 0.257 ± 0.009 | **0.059 ± 0.002** |

Table S13: **NMAE values on major cellular levelpredicted via independent level approach.** Received by comparing the estimated cell proportions with the true cell proportions on the artificial test mixtures. The error bars correspond to ±1 SD obtained over 10 simulation runs. Values below 0.1 are highlighted in bold.

#### 2 Ablation study

| | Ablation model 1 | Ablation model 2 | HIDE |
|---|---|---|---|
| Arterial EC | 0.676 ± 0.024 | 0.618 ± 0.025 | 0.643 ± 0.033 |
| Breast basal cell | **0.884 ± 0.006** | **0.861 ± 0.008** | **0.878 ± 0.005** |
| Breast cancer specific luminal cell | **0.862 ± 0.008** | **0.856 ± 0.01** | **0.907 ± 0.006** |
| Breast cancer specific proliferation luminal cell | 0.601 ± 0.026 | 0.602 ± 0.02 | 0.625 ± 0.02 |
| Capillary EC | 0.687 ± 0.03 | 0.655 ± 0.029 | 0.719 ± 0.022 |
| CD4 T | 0.77 ± 0.01 | 0.734 ± 0.012 | 0.769 ± 0.011 |
| cDC2 | 0.666 ± 0.016 | 0.69 ± 0.013 | 0.689 ± 0.023 |
| CFD fibroblast | 0.796 ± 0.009 | **0.805 ± 0.01** | **0.812 ± 0.01** |
| CXCL1/2/3 fibroblast | **0.891 ± 0.008** | **0.89 ± 0.009** | **0.901 ± 0.008** |
| CXCL13 exhausted CD8 T | 0.54 ± 0.019 | 0.55 ± 0.021 | 0.556 ± 0.015 |
| GZMH CD8 T | 0.588 ± 0.019 | 0.58 ± 0.024 | 0.612 ± 0.022 |
| GZMK CD8 T | 0.757 ± 0.014 | 0.746 ± 0.017 | 0.78 ± 0.015 |
| IgA plasma | **0.888 ± 0.006** | **0.888 ± 0.006** | **0.877 ± 0.007** |
| IgG plasma cell | **0.949 ± 0.004** | **0.937 ± 0.02** | **0.936 ± 0.004** |
| INF responsed T | 0.308 ± 0.035 | 0.27 ± 0.016 | 0.176 ± 0.049 |
| Luminal progenitor | **0.838 ± 0.007** | **0.828 ± 0.007** | **0.838 ± 0.005** |
| Lymphatic EC | **0.814 ± 0.022** | 0.749 ± 0.02 | 0.763 ± 0.017 |
| Macrophage | **0.809 ± 0.015** | **0.828 ± 0.01** | **0.898 ± 0.005** |
| Mast cell | 0.759 ± 0.017 | 0.75 ± 0.016 | 0.745 ± 0.019 |
| Monocyte | 0.358 ± 0.021 | 0.326 ± 0.023 | 0.383 ± 0.018 |
| mregDC | 0.741 ± 0.025 | 0.668 ± 0.022 | 0.651 ± 0.028 |
| NK cell | 0.741 ± 0.012 | 0.755 ± 0.01 | 0.757 ± 0.01 |
| Other B cells | **0.857 ± 0.009** | **0.839 ± 0.01** | **0.84 ± 0.009** |
| Other fibroblasts | **0.903 ± 0.009** | **0.902 ± 0.008** | **0.909 ± 0.007** |
| pDC | **0.919 ± 0.007** | **0.879 ± 0.006** | **0.831 ± 0.017** |
| Pericyte | 0.673 ± 0.016 | 0.617 ± 0.017 | 0.626 ± 0.031 |
| Proliferation macrophage | 0.362 ± 0.026 | 0.379 ± 0.045 | 0.38 ± 0.035 |
| Proliferation T/NK | 0.557 ± 0.025 | 0.574 ± 0.026 | 0.574 ± 0.027 |
| Smooth muscle cell | **0.855 ± 0.01** | **0.827 ± 0.01** | **0.865 ± 0.01** |
| Tfh | 0.749 ± 0.011 | 0.748 ± 0.01 | 0.736 ± 0.01 |
| TGM2 luminal cell | 0.734 ± 0.017 | 0.722 ± 0.014 | 0.735 ± 0.017 |
| TNBC-specific epithelial cell | **0.938 ± 0.006** | **0.934 ± 0.007** | **0.945 ± 0.005** |
| Treg | **0.828 ± 0.012** | **0.814 ± 0.012** | **0.818 ± 0.011** |
| Venous EC | **0.822 ± 0.011** | **0.823 ± 0.009** | **0.839 ± 0.008** |
| mean | 0.739 ± 0.002 | 0.725 ± 0.002 | 0.736 ± 0.002 |

Table S14: **Observed Pearson's correlations on sub cellular level,** by comparing the estimated cell proportions with the true cell proportions predicted in the benchmarking study for artificial test mixtures. The error bars correspond to ±1 SD obtained over 10 simulation runs. Correlations above 0.8 are highlighted in bold.

| | Ablation model 1 | Ablation model 2 | HIDE |
|---|---|---|---|
| B cell | **0.928 ± 0.005** | **0.92 ± 0.007** | **0.931 ± 0.005** |
| CD4 T cell | **0.868 ± 0.008** | **0.817 ± 0.008** | **0.884 ± 0.006** |
| CD8 T cell | 0.783 ± 0.009 | 0.768 ± 0.011 | **0.838 ± 0.006** |
| Dendritic cell | 0.722 ± 0.017 | 0.73 ± 0.015 | 0.754 ± 0.02 |
| Endothelial cell | **0.907 ± 0.005** | **0.897 ± 0.005** | **0.946 ± 0.003** |
| Epithelial cell | **0.947 ± 0.004** | **0.927 ± 0.005** | **0.962 ± 0.003** |
| Fibroblast | **0.945 ± 0.003** | **0.934 ± 0.004** | **0.953 ± 0.003** |
| Granulocyte | 0.759 ± 0.017 | 0.75 ± 0.016 | 0.745 ± 0.019 |
| Macrophage | **0.801 ± 0.013** | **0.868 ± 0.008** | **0.909 ± 0.006** |
| Monocyte | 0.358 ± 0.021 | 0.326 ± 0.023 | 0.383 ± 0.018 |
| NK cell | 0.741 ± 0.011 | 0.755 ± 0.01 | 0.757 ± 0.01 |
| Perivascular cell | **0.891 ± 0.006** | **0.846 ± 0.013** | **0.905 ± 0.006** |
| Proliferation T/NK | 0.555 ± 0.025 | 0.574 ± 0.026 | 0.574 ± 0.027 |
| mean | 0.785 ± 0.004 | 0.778 ± 0.002 | **0.811 ± 0.002** |

Table S15: **Observed Pearson's correlations for the minor cellular level predicted via summing approach.** Pearson's correlations are calculated between true and predited cellular composition. The error bars correspond to ±1 SD obtained over 10 simulation runs. Correlations above 0.8 are highlighted in bold.

| | Ablation model 1 | Ablation model 2 | HIDE |
|---|---|---|---|
| B cell | **0.928 ± 0.005** | **0.92 ± 0.007** | **0.931 ± 0.005** |
| Endothelial cell | **0.907 ± 0.005** | **0.897 ± 0.005** | **0.946 ± 0.003** |
| Epithelial cell | **0.947 ± 0.004** | **0.927 ± 0.005** | **0.962 ± 0.003** |
| Fibroblast | **0.945 ± 0.003** | **0.934 ± 0.004** | **0.953 ± 0.003** |
| Myeloid cell | **0.887 ± 0.007** | **0.754 ± 0.013** | **0.965 ± 0.002** |
| NK cell | 0.741 ± 0.011 | 0.755 ± 0.01 | 0.757 ± 0.01 |
| Perivascular cell | **0.891 ± 0.006** | **0.846 ± 0.013** | **0.905 ± 0.006** |
| Proliferation T/NK | 0.555 ± 0.025 | 0.574 ± 0.026 | 0.574 ± 0.027 |
| T cell | **0.892 ± 0.007** | **0.886 ± 0.006** | **0.924 ± 0.003** |
| mean | **0.855 ± 0.004** | **0.833 ± 0.004** | **0.88 ± 0.003** |

Table S16: **Observed Pearson's correlations for the major cellular level predicted via summing approach.** Pearson's correlations are calculated between true and predited cellular composition. The error bars correspond to ±1 SD obtained over 10 simulation runs. Correlations above 0.8 are highlighted in bold.

| | Ablation model 1 | Ablation model 2 | HIDE |
|---|---|---|---|
| Arterial EC | **0.093 ± 0.012** | 0.25 ± 0.036 | **0.089 ± 0.012** |
| Breast basal cell | **0.07 ± 0.01** | **0.099 ± 0.015** | **0.07 ± 0.01** |
| Breast cancer specific luminal cell | **0.076 ± 0.006** | 0.464 ± 0.036 | **0.053 ± 0.004** |
| Breast cancer specific proliferation luminal cell | 0.153 ± 0.016 | 0.113 ± 0.012 | 0.107 ± 0.01 |
| Capillary EC | 0.113 ± 0.014 | 0.333 ± 0.039 | 0.103 ± 0.012 |
| CD4 T | 0.106 ± 0.01 | 0.117 ± 0.012 | **0.084 ± 0.008** |
| cDC2 | 0.138 ± 0.012 | 0.996 ± 0.097 | 0.118 ± 0.012 |
| CFD fibroblast | **0.095 ± 0.013** | 0.123 ± 0.016 | **0.084 ± 0.011** |
| CXCL1/2/3 fibroblast | **0.064 ± 0.008** | **0.089 ± 0.01** | **0.058 ± 0.007** |
| CXCL13 exhausted CD8 T | 0.163 ± 0.022 | 0.129 ± 0.014 | 0.131 ± 0.014 |
| GZMH CD8 T | 0.161 ± 0.015 | 0.167 ± 0.014 | 0.111 ± 0.01 |
| GZMK CD8 T | 0.106 ± 0.012 | 0.153 ± 0.017 | **0.086 ± 0.01** |
| IgA plasma | **0.052 ± 0.008** | **0.081 ± 0.012** | **0.06 ± 0.009** |
| IgG plasma cell | **0.042 ± 0.005** | 0.152 ± 0.018 | **0.052 ± 0.006** |
| INF responsed T | 0.198 ± 0.032 | 0.883 ± 0.135 | 0.101 ± 0.014 |
| Luminal progenitor | **0.084 ± 0.009** | 0.105 ± 0.009 | **0.076 ± 0.008** |
| Lymphatic EC | **0.067 ± 0.013** | 0.381 ± 0.076 | **0.059 ± 0.011** |
| Macrophage | **0.09 ± 0.005** | **0.098 ± 0.006** | **0.059 ± 0.004** |
| Mast cell | **0.09 ± 0.011** | 0.169 ± 0.026 | **0.094 ± 0.013** |
| Monocyte | 0.231 ± 0.026 | 0.509 ± 0.057 | 0.143 ± 0.018 |
| mregDC | **0.072 ± 0.016** | 0.495 ± 0.098 | **0.058 ± 0.012** |
| NK cell | 0.112 ± 0.012 | 0.15 ± 0.016 | 0.**0.097 ± 0.011** |
| Other B cells | **0.077 ± 0.009** | 0.275 ± 0.038 | **0.082 ± 0.011** |
| Other fibroblasts | **0.062 ± 0.005** | **0.098 ± 0.006** | **0.057 ± 0.004** |
| pDC | **0.043 ± 0.006** | 0.387 ± 0.05 | **0.066 ± 0.01** |
| Pericyte | 0.108 ± 0.017 | 0.424 ± 0.073 | **0.098 ± 0.018** |
| Proliferation macrophage | 0.202 ± 0.017 | 0.461 ± 0.041 | 0.139 ± 0.013 |
| Proliferation T/NK | 0.171 ± 0.019 | 0.139 ± 0.015 | 0.13 ± 0.014 |
| Smooth muscle cell | **0.076 ± 0.01** | 0.309 ± 0.036 | **0.073 ± 0.009** |
| Tfh | 0.115 ± 0.014 | 0.104 ± 0.012 | **0.099 ± 0.012** |
| TGM2 luminal cell | 0.101 ± 0.013 | **0.097 ± 0.013** | **0.093 ± 0.012** |
| TNBC-specific epithelial cell | **0.045 ± 0.004** | 0.187 ± 0.016 | **0.04 ± 0.004** |
| Treg | **0.088 ± 0.006** | 0.102 ± 0.007 | **0.079 ± 0.006** |
| Venous EC | **0.083 ± 0.014** | 0.118 ± 0.017 | **0.079 ± 0.012** |
| mean | 0.104 ± 0.003 | 0.258 ± 0.007 | **0.086 ± 0.002** |

Table S17: **NMAE error on sub cellular level.** Received by comparing the estimated cell proportions with the true cell proportions on the artificial test mixtures. The error bars correspond to ±1 SD obtained over 10 simulation runs. Values below 0.1 are highlighted in bold.

| | Ablation model 1 | Ablation model 2 | HIDE |
|---|---|---|---|
| B cell | **0.05 ± 0.004** | 0.244 ± 0.034 | **0.047 ± 0.004** |
| CD4 T cell | **0.072 ± 0.005** | 0.391 ± 0.034 | **0.058 ± 0.004** |
| CD8 T cell | **0.095 ± 0.007** | 0.247 ± 0.009 | **0.069 ± 0.005** |
| Dendritic cell | 0.126 ± 0.017 | 2.229 ± 0.212 | **0.099 ± 0.015** |
| Endothelial cell | **0.07 ± 0.008** | 0.745 ± 0.047 | **0.046 ± 0.005** |
| Epithelial cell | **0.045 ± 0.004** | 1.821 ± 0.126 | **0.035 ± 0.003** |
| Fibroblast | **0.043 ± 0.003** | 0.305 ± 0.023 | **0.038 ± 0.003** |
| Granulocyte | **0.094 ± 0.012** | 0.254 ± 0.036 | **0.094 ± 0.013** |
| Macrophage | 0.1 ± 0.008 | 0.269 ± 0.024 | **0.054 ± 0.004** |
| Monocyte | 0.239 ± 0.027 | 0.421 ± 0.034 | 0.143 ± 0.018 |
| NK cell | 0.116 ± 0.012 | 0.314 ± 0.044 | **0.097 ± 0.011** |
| Perivascular cell | **0.072 ± 0.008** | 1.088 ± 0.082 | **0.061 ± 0.007** |
| Proliferation T/NK | 0.178 ± 0.02 | 0.243 ± 0.029 | 0.13 ± 0.014 |
| mean | 0.1 ± 0.003 | 0.659 ± 0.026 | **0.075 ± 0.003** |

Table S18: **NMAE error on minor cellular level predicted via summing approach.** Received by comparing the estimated cell proportions with the true cell proportions on the artificial test mixtures. The error bars correspond to ±1 SD obtained over 10 simulation runs. Values below 0.1 are highlighted in bold.

| | Ablation model 1 | Ablation model 2 | HIDE |
|---|---|---|---|
| B cell | **0.05 ± 0.004** | 0.244 ± 0.034 | **0.047 ± 0.004** |
| Endothelial cell | **0.07 ± 0.008** | 0.745 ± 0.047 | **0.046 ± 0.005** |
| Epithelial cell | **0.045 ± 0.004** | 1.821 ± 0.126 | **0.035 ± 0.003** |
| Fibroblast | **0.043 ± 0.003** | 0.305 ± 0.023 | **0.038 ± 0.003** |
| Myeloid cell | **0.088 ± 0.008** | 1.293 ± 0.085 | **0.033 ± 0.003** |
| NK cell | 0.116 ± 0.012 | 0.314 ± 0.044 | **0.097 ± 0.011** |
| Perivascular cell | **0.072 ± 0.008** | 1.088 ± 0.082 | **0.061 ± 0.007** |
| Proliferation T/NK | 0.178 ± 0.02 | 0.243 ± 0.029 | 0.13 ± 0.014 |
| T cell | **0.066 ± 0.004** | 0.201 ± 0.011 | **0.047 ± 0.003** |
| mean | **0.081 ± 0.003** | 0.695 ± 0.028 | **0.059 ± 0.002** |

Table S19: **NMAE error on major cellular level predicted via summing approach.** Received by comparing the estimated cell proportions with the true cell proportions on the artificial test mixtures. The error bars correspond to ±1 SD obtained over 10 simulation runs. Values below 0.1 are highlighted in bold.

### 3 TCGA HIDE

| | all subtypes | HER2 | LumA | LumB | TNBC |
|---|---|---|---|---|---|
| B cell | -12.69 | -15.16 | 5.46 | -6.73 | -5.85 |
| Endothelial cell | 5.98 | 6.25 | 0.96 | -2.45 | 1.29 |
| Epithelial cell | 0.58 | **7.04** | -0.01 | 0.49 | 0.68 |
| Fibroblast | 0.33 | **-9.20** | 0.44 | **5.36** | 0.38 |
| Myeloid cell | -1.76 | **-16.55** | -5.15 | 1.28 | -3.17 |
| NK cell | -11.65 | **-41.99** | -15.93 | **-37.69** | **-21.05** |
| Perivascular cell | 4.34 | 3.65 | **8.65** | -2.39 | **6.13** |
| Proliferation T/NK | -15.45 | 3.99 | -6.23 | 1.09 | -4.01 |
| T cell | 0.21 | -7.10 | -1.62 | -2.85 | -1.69 |

Table S20: **Coefficients of the Cox proportional hazard model** for all major celltypes. Coefficients having a p-value smaller than 0.05 are highlighted in bold.

| | all subtypes | HER2 | LumA | LumB | TNBC |
|---|---|---|---|---|---|
| B cell | 0.32 | 0.35 | 0.56 | 0.54 | 0.33 |
| Endothelial cell | 0.26 | 0.43 | 0.65 | 0.55 | 0.45 |
| Epithelial cell | 0.74 | **0.0015** | 0.99 | 0.68 | 0.33 |
| Fibroblast | 0.91 | **0.047** | 0.86 | **0.033** | 0.79 |
| Myeloid cell | 0.66 | **0.0095** | 0.24 | 0.77 | 0.19 |
| NK cell | 0.07 | **0.037** | 0.19 | **0.02** | **0.0013** |
| Perivascular cell | 0.59 | 0.81 | **1.97e-06** | 0.69 | **0.013** |
| Proliferation T/NK | 0.34 | 0.75 | 0.72 | 0.86 | 0.50 |
| T cell | 0.91 | 0.14 | 0.43 | 0.14 | 0.14 |

Table S21: **p-values of the Cox proportional hazard model** for all major celltypes. p-Values below 0.05 are highlighted in bold.

| | all subtypes | HER2 | LumA | LumB | TNBC |
|---|---|---|---|---|---|
| B cell | -12.69 | -15.16 | 5.46 | -6.73 | -5.85 |
| CD4 T cell | 7.30 | -3.59 | **-10.10** | -0.95 | -1.72 |
| CD8 T cell | -1.53 | -8.09 | 0.46 | -3.82 | -1.59 |
| Dendritic cell | **-32.16** | -16.87 | -18.52 | -17.40 | **-23.16** |
| Endothelial cell | 5.98 | 6.25 | 0.96 | -2.45 | 1.29 |
| Epithelial cell | 0.58 | **7.04** | -0.01 | 0.49 | 0.68 |
| Fibroblast | 0.33 | **-9.20** | 0.44 | **5.36** | 0.38 |
| Granulocyte | 34.17 | **-146.24** | -71.30 | 85.01 | -14.90 |
| Macrophage | 0.17 | **-20.31** | -5.08 | 4.64 | -1.48 |
| Monocyte | 7.20 | -240.64 | -66.99 | -101.17 | -20.30 |
| NK cell | -11.65 | **-41.99** | -15.93 | **-37.69** | **-21.05** |
| Perivascular cell | 4.34 | 3.65 | **8.65** | -2.39 | **6.13** |
| Proliferation T/NK | -15.45 | 3.99 | -6.23 | 1.09 | -4.01 |

Table S22: **Coefficients of the Cox proportional hazard model** for all minor celltypes. Coefficients having a p-Value below 0.05 are highlighted in bold.

| | all subtypes | HER2 | LumA | LumB | TNBC |
|---|---|---|---|---|---|
| B cell | 0.32 | 0.35 | 0.56 | 0.54 | 0.33 |
| CD4 T cell | 0.053 | 0.64 | **0.023** | 0.86 | 0.54 |
| CD8 T cell | 0.47 | 0.12 | 0.78 | 0.15 | 0.18 |
| Dendritic cell | **0.017** | 0.48 | 0.18 | 0.24 | **0.0031** |
| Endothelial cell | 0.26 | 0.43 | 0.65 | 0.55 | 0.45 |
| Epithelial cell | 0.74 | **0.0015** | 0.99 | 0.68 | 0.33 |
| Fibroblast | 0.91 | **0.047** | 0.86 | **0.033** | 0.79 |
| Granulocyte | 0.54 | **0.024** | 0.14 | 0.17 | 0.62 |
| Macrophage | 0.97 | **0.033** | 0.43 | 0.46 | 0.64 |
| Monocyte | 0.72 | 0.071 | 0.40 | 0.07 | 0.34 |
| NK cell | 0.07 | **0.037** | 0.19 | **0.02** | **0.0013** |
| Perivascular cell | 0.59 | 0.81 | **1.97e-06** | 0.69 | **0.013** |
| Proliferation T/NK | 0.34 | 0.75 | 0.72 | 0.86 | 0.50 |

Table S23: **p-values of the Cox proportional hazard model** for all minor celltypes.

p-Values below 0.05 are highlighted in bold.

| | all subtypes | HER2 | LumA | LumB | TNBC |
|---|---|---|---|---|---|
| Arterial EC | -101.28 | **43.41** | 8.79 | 33.84 | 16.06 |
| Breast basal cell | -8.48 | 14.53 | 8.91 | **41.57** | 4.17 |
| Breast cancer specific luminal cell | **6.67** | **7.54** | 0.43 (0.69) | -0.41 | 1.03 |
| Breast cancer specific proliferation luminal cell | -0.11 | 3.92 | -3.29 | 3.93 | 1.76 |
| Capillary EC | 4.42 | 2.76 | 1.54 | -7.10 | 0.96 |
| CD4 T | 5.44 | -5.71 | **-9.56** | -1.71 | -3.01 |
| cDC2 | -34.92 | -34.29 | -19.38 | -4.35 | **-22.78** |
| CFD fibroblast | 3.21 | -12.39 | 0.36 | 5.26 | 0.63 |
| CXCL1/2/3 fibroblast | -37.48 | -12.79 | -18.85 | 17.57 | -12.60 |
| CXCL13 exhausted CD8 T | **-39.82** | -192.43 | **-123.68** | -30.21 | **-45.53** |
| GZMH CD8 T | 3.40 | -11.75 | 1.29 | -7.33 | -0.14 |
| GZMK CD8 T | -2.68 | -14.09 | 0.76 | -6.35 | -2.91 |
| IgA plasma | -260.41 | **-949.51** | 179.55 | 18.35 | -86.64 |
| IgG plasma cell | -124.48 | -369.38 | 3.53 | -2.17 | -77.70 |
| INF responsed T | **80.28** | -5602.28 | -60.85 | 20.41 | 33.68 |
| Luminal progenitor | -1.17 | -4.92 | **-31.29** | -29.94 | -7.14 |
| Lymphatic EC | 15.86 | -3.06 | -18.65 | 30.54 | 0.38 |
| Macrophage | -1.67 | **-21.08** | -3.37 | 2.36 | -2.45 |
| Mast cell | 34.17 | **-146.24** | -71.30 | 85.01 | -14.90 |
| Monocyte | 7.20 | -240.64 | -66.99 | -101.17 | -20.30 |
| mregDC | **-93.12** | 22.12 | -67.10 | **-193.46** | **-83.75** |
| NK cell | -11.65 | **-41.99** | -15.93 | **-37.69** | **-21.05** |
| Other B cells | -12.39 | -11.35 | 5.76 | -8.51 | -5.54 |
| Other fibroblasts | -0.42 | -15.39 | 2.25 | **11.33** | 2.24 |
| pDC | 45.06 | 23.47 | -30.35 | -91.80 | -26.27 |
| Pericyte | -8.50 | 44.12 | **7.72** | 15.86 | **7.89** |
| Proliferation macrophage | 31.82 | -17.72 | -53.75 | 32.55 | 13.24 |
| Proliferation T/NK | -15.45 | 3.99 | -6.23 | 1.09 | -4.01 |
| Smooth muscle cell | 5.88 | -14.89 | 11.25 | -6.99 | 3.94 |
| Tfh | 30.05 | -12.58 | -75.13 | **-160.34** | -62.06 |
| TGM2 luminal cell | -11.76 | 8.05 | -0.69 | 2.33 | 0.69 |
| TNBC-specific epithelial cell | -2.55 | -3.38 | -19.74 | **16.79** | -1.99 |
| Treg | 62.85 | 62.94 | 5.82 | 43.95 | **27.90** |
| Venous EC | 4.65 | 11.23 | -20.41 | **24.87** | -0.52 |

Table S24: **Coefficients of the Cox proportional hazard model** for all sub celltypes. Coefficients having a p-Value below 0.05 are highlighted in bold.

| | all subtypes | HER2 | LumA | LumB | TNBC |
|---|---|---|---|---|---|
| Arterial EC | 0.20 | **0.020** | 0.61 | 0.16 | 0.18 |
| Breast basal cell | 0.49 | 0.60 | 0.20 | **0.035** | 0.46 |
| Breast cancer specific luminal cell | **0.04** | **0.0066** | 0.69 | 0.71 | 0.19 |
| Breast cancer specific proliferation luminal cell | 0.98 | 0.51 | 0.65 | 0.16 | 0.40 |
| Capillary EC | 0.34 | 0.78 | 0.46 | 0.23 | 0.59 |
| CD4 T | 0.25 | 0.49 | **0.026** | 0.79 | 0.31 |
| cDC2 | 0.07 | 0.25 | 0.22 | 0.84 | **0.025** |
| CFD fibroblast | 0.48 | 0.11 | 0.89 | 0.21 | 0.74 |
| CXCL1/2/3 fibroblast | 0.13 | 0.10 | 0.41 | 0.29 | 0.10 |
| CXCL13 exhausted CD8 T | **0.0036** | 0.31 | **0.036** | 0.15 | **0.00072** |
| GZMH CD8 T | 0.41 | 0.20 | 0.61 | 0.25 | 0.95 |
| GZMK CD8 T | 0.61 | 0.26 | 0.84 | 0.21 | 0.29 |
| IgA plasma | 0.33 | **0.037** | 0.36 | 0.93 | 0.59 |
| IgG plasma cell | 0.24 | 0.06 | 0.97 | 0.98 | 0.18 |
| INF responsed T | **0.00028** | 0.23 | 0.33 | 0.72 | 0.11 |
| Luminal progenitor | 0.80 | 0.58 | **0.0021** | 0.26 | 0.11 |
| Lymphatic EC | 0.49 | 0.92 | 0.23 | 0.22 | 0.97 |
| Macrophage | 0.73 | **0.047** | 0.57 | 0.73 | 0.45 |
| Mast cell | 0.54 | **0.024** | 0.14 | 0.17 | 0.62 |
| Monocyte | 0.72 | 0.07 | 0.40 | 0.07 | 0.34 |
| mregDC | **0.018** | 0.72 | 0.40 | **0.037** | **0.0037** |
| NK cell | 0.073 | **0.037** | 0.19 | **0.020** | **0.0013** |
| Other B cells | 0.37 | 0.56 | 0.56 | 0.48 | 0.41 |
| Other fibroblasts | 0.94 | 0.17 | 0.71 | **0.023** | 0.47 |
| pDC | 0.57 | 0.86 | 0.13 | 0.20 | 0.48 |
| Pericyte | 0.79 | 0.15 | **7.63e-16** | 0.42 | **1.5e-13** |
| Proliferation macrophage | 0.14 | 0.53 | 0.15 | 0.13 | 0.33 |
| Proliferation T/NK | 0.34 | 0.75 | 0.72 | 0.86 | 0.50 |
| Smooth muscle cell | 0.48 | 0.45 | 0.15 | 0.42 | 0.38 |
| Tfh | 0.65 | 0.87 | 0.29 | **0.03** | 0.15 |
| TGM2 luminal cell | 0.48 | 0.35 | 0.92 | 0.82 | 0.88 |
| TNBC-specific epithelial cell | 0.51 | 0.68 | 0.27 | **0.03** | 0.55 |
| Treg | 0.093 | 0.29 | 0.77 | 0.18 | **0.049** |
| Venous EC | 0.86 | 0.53 | 0.13 | **0.04** | 0.95 |

Table S25: **p-values of the Cox proportional hazard model** for all sub celltypes. p-Values below 0.05 are highlighted in bold.

| celltype | T>0N0M0 | T>0N>0M>0 | p |
|---|---|---|---|
| Endothelial cell | 0.1085 ± 0.0493 | 0.1029 ± 0.0456 | 0.3048 |
| Epithelial cell | 0.4275 ± 0.1289 | 0.4192 ± 0.1342 | 0.7495 |
| Fibroblast | 0.0954 ± 0.0588 | 0.0937 ± 0.0602 | 0.6406 |
| B cell | 0.0268 ± 0.0155 | 0.0314 ± 0.0176 | **0.0059** |
| NK cell | 0.0334 ± 0.0136 | 0.0331 ± 0.0219 | **0.0354** |
| Proliferation T NK | 0.0115 ± 0.0133 | 0.0156 ± 0.0169 | **0.0449** |
| T cell | 0.1892 ± 0.0648 | 0.1910 ± 0.0765 | 0.8052 |
| Myeloid cell | 0.0711 ± 0.0354 | 0.0752 ± 0.0447 | 0.7183 |
| Perivascular cell | 0.0366 ± 0.0225 | 0.0378 ± 0.0277 | 0.9869 |
| CD4 T cell | 0.0763 ± 0.0325 | 0.0742 ± 0.0341 | 0.5799 |
| CD8 T cell | 0.1129 ± 0.0620 | 0.1167 ± 0.0717 | 0.9467 |
| CD4 T | 0.0625 ± 0.0304 | 0.0613 ± 0.0310 | 0.7341 |
| Tfh | 0.0029 ± 0.0037 | 0.0026 ± 0.0052 | 0.3138 |
| Treg | 0.0091 ± 0.0049 | 0.0090 ± 0.0054 | 0.6891 |
| INF responsed T | 0.0019 ± 0.0047 | 0.0014 ± 0.0036 | 0.8225 |
| CXCL13 exhausted CD8 T | 0.0049 ± 0.0116 | 0.0072 ± 0.0227 | 0.5145 |
| GZMH CD8 T | 0.0776 ± 0.0351 | 0.0781 ± 0.0370 | 0.8780 |
| GZMK CD8 T | 0.0305 ± 0.0291 | 0.0314 ± 0.0353 | 0.5732 |
| other B cells | 0.0241 ± 0.0137 | 0.0288 ± 0.0164 | **0.0048** |
| IgA plasma | 0.0007 ± 0.0009 | 0.0006 ± 0.0007 | 0.3838 |
| IgG plasma cell | 0.0020 ± 0.0025 | 0.0020 ± 0.0016 | **0.0334** |
| Arterial EC | 0.0047 ± 0.0075 | 0.0030 ± 0.0053 | **0.0009** |
| Capillary EC | 0.0843 ± 0.0506 | 0.0843 ± 0.0486 | 0.7470 |
| Lymphatic EC | 0.0091 ± 0.0069 | 0.0068 ± 0.0066 | **0.0002** |
| Venous EC | 0.0104 ± 0.0120 | 0.0087 ± 0.0090 | 0.2689 |
| Luminal progenitor | 0.0179 ± 0.0255 | 0.0264 ± 0.0397 | 0.1838 |
| TGM2 luminal cell | 0.0144 ± 0.0156 | 0.0165 ± 0.0173 | 0.3480 |
| Breast cancer specific luminal cell | 0.3145 ± 0.1435 | 0.2785 ± 0.1630 | **0.0497** |
| Breast cancer specific proliferation luminal cell | 0.0427 ± 0.0422 | 0.0489 ± 0.0478 | 0.4173 |
| TNBC-specific epithelial cell | 0.0231 ± 0.0412 | 0.0344 ± 0.0492 | 0.1013 |
| Breast basal cell | 0.0147 ± 0.0133 | 0.0145 ± 0.0122 | 0.8436 |
| Pericyte | 0.0084 ± 0.0081 | 0.0102 ± 0.0176 | 0.4703 |
| Smooth muscle cell | 0.0282 ± 0.0189 | 0.0277 ± 0.0189 | 0.6247 |
| other fibroblasts | 0.0228 ± 0.0251 | 0.0233 ± 0.0244 | 0.5831 |
| CFD fibroblast | 0.0669 ± 0.0497 | 0.0614 ± 0.0462 | 0.2686 |
| CXCL1/2/3 fibroblast | 0.0057 ± 0.0085 | 0.0090 ± 0.0219 | **0.0038** |
| Dendritic cell | 0.0172 ± 0.0093 | 0.0187 ± 0.0122 | 0.4760 |
| Granulocyte | 0.0071 ± 0.0034 | 0.0070 ± 0.0037 | 0.6150 |
| Macrophages | 0.0421 ± 0.0259 | 0.0447 ± 0.0324 | 0.9269 |
| Monocyte | 0.0047 ± 0.0086 | 0.0049 ± 0.0078 | 0.4500 |
| Macrophage | 0.0342 ± 0.0244 | 0.0359 ± 0.0302 | 0.7008 |
| Proliferation macrophage | 0.0079 ± 0.0075 | 0.0088 ± 0.0075 | 0.1769 |
| cDC2 | 0.0140 ± 0.0082 | 0.0147 ± 0.0094 | 0.6507 |
| mregDC | 0.0019 ± 0.0036 | 0.0023 ± 0.0046 | 0.5732 |
| pDC | 0.0013 ± 0.0015 | 0.0018 ± 0.0022 | **0.0387** |

Table S26: **Kruskal Wallis Test for patients with only a primary tumor and patients having lymphnode involvement and metastatis.** Unknown Lymphnode involvement (NX) and metastasis levels (MX) are regarded as present. p-Values below 0.05 are highlighted in bold.

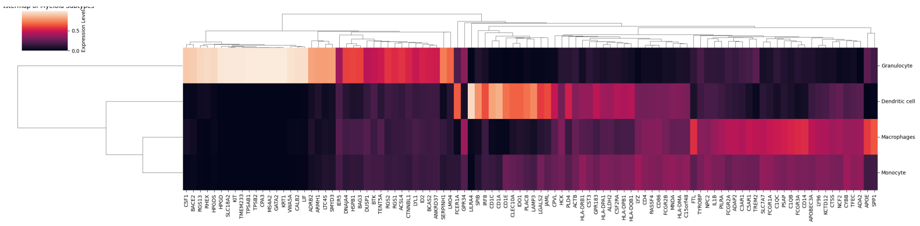


Figure S1: **Heatmap of the 50 most important genes** calculated via the HIDE algorithm. Genes were ranked by $g_{Myelo.} \times var(X_{Myelo,.)})$. Black corresponds to low expression and bright to high expression.

### 4 TCGA Bayesprism

| celltype | All | Her2 | LumA | LumB | TNBC |
|---|---|---|---|---|---|
| B cell | -9.4026 | **-64.7146** | -14.1094 | -0.4991 | -13.2918 |
| Endothelial cell | 0.3126 | 0.2795 | 0.9111 | -3.8546 | 0.7176 |
| Epithelial cell | 0.7337 | **4.1682** | 0.3881 | -0.0016 | 0.9455 |
| Fibroblast | 0.2855 | **-11.3337** | -0.6728 | **5.5736** | 1.8163 |
| Myeloid cell | -1.3378 | -7.4548 | -2.7269 | 0.0410 | 0.0638 |
| NK cell | -3.6520 | -17.1880 | -2.4874 | -4.5490 | **-6.5682** |
| Perivascular cell | 1.5931 | -3.7654 | 0.7040 | 7.4556 | 0.8324 |
| Proliferation T/NK | -0.4208 | -2.8571 | 0.4449 | 2.2205 | -3.9174 |
| T cell | -3.2064 | 3.7769 | -0.1299 | **-8.5666** | **6.9383** |

Table S27: **Coefficients of the Cox proportional hazard model** for all Bayesprism major celltypes using the independent level approach. Coefficients having a p-Value below 0.05 are highlighted in bold.

| celltype | All | Her2 | LumA | LumB | TNBC |
|---|---|---|---|---|---|
| B cell | 0.0739 | **0.0247** | 0.2727 | 0.9425 | 0.1384 |
| Endothelial cell | 0.8061 | 0.9434 | 0.5658 | 0.3066 | 0.8248 |
| Epithelial cell | 0.1839 | **0.0005** | 0.6267 | 0.9987 | 0.4969 |
| Fibroblast | 0.8546 | **0.0471** | 0.7792 | **0.0250** | 0.4165 |
| Myeloid cell | 0.3675 | 0.0740 | 0.2624 | 0.9878 | 0.9803 |
| NK cell | 0.0827 | 0.0957 | 0.3030 | 0.3337 | **0.0256** |
| Perivascular cell | 0.3947 | 0.5298 | 0.7756 | 0.1593 | 0.8000 |
| Proliferation T/NK | 0.7268 | 0.4276 | 0.8404 | 0.2113 | 0.1371 |
| T cell | 0.1294 | 0.4712 | 0.9559 | **0.0303** | **0.0003** |

Table S28: **p-values of the Cox proportional hazard model** for all Bayesprism major using the independent level approach. p-Values below 0.05 are highlighted in bold.

| celltype | All | Her2 | LumA | LumB | TNBC |
|---|---|---|---|---|---|
| B cell | -1.9036 | 1.9999 | 3.2306 | -0.2849 | -3.7519 |
| Endothelial cell | 1.5754 | 1.9921 | 1.2668 | 3.7207 | 1.3504 |
| Epithelial cell | 0.7662 | **3.7228** | 0.4464 | -0.0510 | 0.6071 |
| Fibroblast | 0.3598 | -9.9653 | -0.9926 | 4.9110 | 3.1663 |
| Myeloid cell | -2.0891 | **-9.5628** | -3.7460 | 0.2339 | 0.1495 |
| NK cell | -27.4687 | -12.4174 | -42.2026 | -33.3553 | 9.3520 |
| Perivascular cell | 5.3841 | **-26.2962** | **13.6436** | 7.3365 | -3.1559 |
| Proliferation T/NK | -2.8569 | -6.3572 | 3.4119 | -7.9954 | -1.9389 |
| T cell | -3.6986 | -7.8742 | -1.3963 | -5.7811 | -3.1066 |

Table S29: **Coefficients of the Cox proportional hazard model** for all Bayesprism major celltypes using the summing approach. Coefficients having a p-Value below 0.05 are highlighted in bold.

| celltype | All | Her2 | LumA | LumB | TNBC |
|---|---|---|---|---|---|
| B cell | 0.3045 | 0.6279 | 0.3980 | 0.9114 | 0.3697 |
| Endothelial cell | 0.1326 | 0.7043 | 0.4668 | 0.2881 | 0.3186 |
| Epithelial cell | 0.1827 | **0.0019** | 0.6045 | 0.9624 | 0.6660 |
| Fibroblast | 0.8046 | 0.1167 | 0.5965 | 0.0868 | 0.3753 |
| Myeloid cell | 0.1192 | **0.0051** | 0.0654 | 0.9264 | 0.9501 |
| NK cell | 0.0631 | 0.5745 | 0.0548 | 0.5503 | 0.4839 |
| Perivascular cell | 0.1444 | **0.0266** | **0.0002** | 0.4229 | 0.5474 |
| Proliferation T/NK | 0.2602 | 0.3242 | 0.6601 | 0.2851 | 0.6005 |
| T cell | 0.0179 | 0.0738 | 0.5718 | 0.0597 | 0.1789 |

Table S30: **p-values of the Cox proportional hazard model** for all Bayesprism major using the summing approach. p-Values below 0.05 are highlighted in bold.

| celltype | All | Her2 | LumA | LumB | TNBC |
|---|---|---|---|---|---|
| B cell | -8.3739 | **-61.9155** | -12.5668 | -0.5659 | -12.8468 |
| CD4 T cell | -0.9286 | 10.2179 | 0.9062 | -2.4928 | **10.1101** |
| CD8 T cell | **-5.8444** | -21.1047 | -3.1365 | -9.7985 | -6.5525 |
| Dendritic cell | 0.9245 | 3.5107 | 2.2533 | 1.0173 | -4.0091 |
| Endothelial cell | 0.4199 | 0.3037 | 0.9572 | -4.1710 | 0.9648 |
| Epithelial cell | 0.8298 | **3.8595** | 0.4985 | -0.1420 | 0.6830 |
| Fibroblast | 0.2646 | **-10.9716** | -0.3389 | **5.1962** | 0.8777 |
| Granulocyte | -0.5295 | 2.0843 | -2.4196 | **8.1836** | **4.0152** |
| Macrophage | 0.0882 | -3.8284 | -3.1627 | 1.9051 | 2.5938 |
| Monocyte | -2.3031 | -5.6797 | -1.7471 | -3.2309 | -1.6619 |
| NK cell | 0.9951 | -19.8187 | -1.8088 | -1.5835 | **12.1541** |
| Perivascular cell | 2.9622 | -7.5198 | 6.8891 | 2.3795 | -1.0588 |
| Proliferation T/NK | -0.9049 | -0.9283 | 1.6681 | 0.9201 | -3.0189 |

Table S31: **Coefficients of the Cox proportional hazard model** for all bayesprism minor celltypes using the independent level approach. Coefficient having a p-Value below 0.05 are highlighted in bold.

| celltype | All | Her2 | LumA | LumB | TNBC |
|---|---|---|---|---|---|
| B cell | 0.0704 | **0.0279** | 0.2695 | 0.9265 | 0.1352 |
| CD4 T cell | 0.7032 | 0.1509 | 0.7540 | 0.5626 | **0.0308** |
| CD8 T cell | **0.0309** | 0.0905 | 0.3650 | 0.0757 | 0.0510 |
| Dendritic cell | 0.5993 | 0.5070 | 0.2801 | 0.8156 | 0.2999 |
| Endothelial cell | 0.7380 | 0.9417 | 0.5430 | 0.3175 | 0.7541 |
| Epithelial cell | 0.1484 | **0.0010** | 0.5371 | 0.8983 | 0.6530 |
| Fibroblast | 0.8660 | **0.0427** | 0.8901 | **0.0315** | 0.7414 |
| Granulocyte | 0.6643 | 0.6467 | 0.1408 | **0.0037** | **0.0115** |
| Macrophage | 0.9599 | 0.5234 | 0.2603 | 0.6164 | 0.3963 |
| Monocyte | 0.1589 | 0.2579 | 0.5375 | 0.2737 | 0.4894 |
| NK cell | 0.7677 | 0.2156 | 0.6223 | 0.8377 | **0.0037** |
| Perivascular cell | 0.2330 | 0.3811 | 0.0729 | 0.6848 | 0.7756 |
| Proliferation T/NK | 0.5034 | 0.8023 | 0.4889 | 0.6604 | 0.2718 |

Table S32: **p-values of the Cox proportional hazard model** for all bayesprism minor celltypes using the independent level approach. p-Values below 0.05 are highlighted in bold.

| celltype | All | Her2 | LumA | LumB | TNBC |
|---|---|---|---|---|---|
| B cell | -1.9036 | 1.9999 | 3.2306 | -0.2849 | -3.7519 |
| CD4 T cell | 1.5606 | 3.0184 | 0.2464 | 0.0122 | 7.1549 |
| CD8 T cell | **-4.5455** | -12.5079 | -1.3808 | -5.4063 | **-6.7251** |
| Dendritic cell | -4.2210 | -18.3559 | -2.4806 | -7.2820 | -6.6139 |
| Endothelial cell | 1.5754 | 1.9921 | 1.2668 | 3.7207 | 1.3504 |
| Epithelial cell | 0.7662 | **3.7228** | 0.4464 | -0.0510 | 0.6071 |
| Fibroblast | 0.3598 | -9.9653 | -0.9926 | 4.9110 | 3.1663 |
| Granulocyte | -0.6795 | -18.4340 | -4.3766 | **9.6800** | 7.9100 |
| Macrophage | 1.3040 | -1.7956 | -3.1875 | 4.9482 | 3.9999 |
| Monocyte | **-5.9500** | -5.5658 | 0.6082 | **-10.2672** | -4.0897 |
| NK cell | -27.4687 | -12.4174 | -42.2026 | -33.3553 | 9.3520 |
| Perivascular cell | 5.3841 | **-26.2962** | **13.6436** | 7.3365 | -3.1559 |
| Proliferation T/NK | -2.8569 | -6.3572 | 3.4119 | -7.9954 | -1.9389 |

Table S33: **Coefficients of the Cox proportional hazard model** for all bayesprism minor celltypes using the summing approach. Coefficients having a p-Value below 0.05 are highlighted in bold.

| celltype | All | Her2 | LumA | LumB | TNBC |
|---|---|---|---|---|---|
| B cell | 0.3045 | 0.6279 | 0.3980 | 0.9114 | 0.3697 |
| CD4 T cell | 0.4935 | 0.5952 | 0.9372 | 0.9974 | 0.0817 |
| CD8 T cell | **0.0114** | 0.1216 | 0.5559 | 0.0874 | **0.0434** |
| Dendritic cell | 0.1202 | 0.1479 | 0.4813 | 0.2911 | 0.3735 |
| Endothelial cell | 0.1326 | 0.7043 | 0.4668 | 0.2881 | 0.3186 |
| Epithelial cell | 0.1827 | **0.0019** | 0.6045 | 0.9624 | 0.6660 |
| Fibroblast | 0.8046 | 0.1167 | 0.5965 | 0.0868 | 0.3753 |
| Granulocyte | 0.8139 | 0.4482 | 0.2337 | **0.0009** | 0.3527 |
| Macrophage | 0.3569 | 0.6384 | 0.2017 | 0.0962 | 0.1079 |
| Monocyte | **0.0238** | 0.3585 | 0.8823 | **0.0364** | 0.2143 |
| NK cell | 0.0631 | 0.5745 | 0.0548 | 0.5503 | 0.4839 |
| Perivascular cell | 0.1444 | **0.0266** | **0.0002** | 0.4229 | 0.5474 |
| Proliferation T/NK | 0.2602 | 0.3242 | 0.6601 | 0.2851 | 0.6005 |

Table S34: **p-values of the Cox proportional hazard model** for all bayesprism minor celltypes using the summing approach. p-Values below 0.05 are highlighted in bold.

| celltype | All | Her2 | LumA | LumB | TNBC |
|---|---|---|---|---|---|
| Arterial EC | **5.4162** | 5.2465 | 4.1931 | **9.2239** | 2.3352 |
| Breast basal cell | 3.7439 | -87.3063 | -6.2313 | 28.6982 | 4.7195 |
| Breast cancer specific luminal cell | 0.6825 | 1.0364 | 1.5076 | 0.1862 | 2.4618 |
| Breast cancer specific proliferation luminal cell | 0.8133 | 1.0754 | -0.8467 | -0.3814 | -1.1846 |
| Capillary EC | -0.0650 | 3.8104 | 3.2469 | **-17.3285** | 0.6174 |
| CD4 T | 4.5263 | **18.6437** | 3.0040 | 3.3209 | 4.7720 |
| cDC2 | -2.4607 | -6.7605 | 0.1389 | -21.1442 | -11.7422 |
| CFD fibroblast | -0.5641 | -6.7215 | -1.8336 | 0.5398 | 4.5876 |
| CXCL1/2/3 fibroblast | -17.4739 | **-346.9702** | -0.4755 | 11.7649 | -6.4773 |
| CXCL13 exhausted CD8 T | **-19.1281** | -29.0025 | -19.9742 | -11.1058 | **-21.5443** |
| GZMH CD8 T | -9.6149 | -57.0463 | -16.7306 | 1.5366 | -16.8151 |
| GZMK CD8 T | -1.6261 | -3.2986 | 0.1871 | -11.2952 | 2.9192 |
| IgA plasma | -10.9764 | **-898.1144** | -6.7250 | 18.6267 | -56.9004 |
| IgG plasma cell | -17.1157 | **-1833.3516** | -38.0984 | -3.1572 | -86.8569 |
| INF responsed T | -2.2890 | -8.7228 | 1.6581 | -2.0449 | -3.1449 |
| Luminal progenitor | -1.4996 | 3.7748 | -4.0715 | -14.2509 | -0.7606 |
| Lymphatic EC | -0.3698 | -8.7084 | -6.1128 | 14.7250 | 0.8503 |
| Macrophage | -2.5452 | -3.6832 | -5.5721 | -2.3108 | -7.0242 |
| Mast cell | -0.6795 | -18.4340 | -4.3766 | **9.6800** | 7.9100 |
| Monocyte | **-5.9500** | -5.5658 | 0.6082 | -10.2672 | -4.0897 |
| mregDC | -5.8331 | **-88.7533** | -10.3124 | 3.8403 | **-32.9024** |
| NK cell | -27.4687 | -12.4174 | -42.2026 | -33.3553 | 9.3520 |
| Other B cells | 0.3459 | 2.9760 | 5.6792 | -0.2962 | -3.2714 |
| Other fibroblasts | 3.2253 | -20.0772 | 2.9833 | 7.5250 | 3.3702 |
| pDC | -1.0041 | -124.1435 | -1.0385 | -58.3741 | **14.9171** |
| Pericyte | 5.4178 | **-475.7124** | **12.4094** | **32.3900** | -13.7003 |
| Proliferation macrophage | 2.5757 | 0.4828 | -1.5789 | **8.0697** | **5.4180** |
| Proliferation T/NK | -2.8569 | -6.3572 | 3.4119 | -7.9954 | -1.9389 |
| Smooth muscle cell | 4.2939 | -13.4208 | 10.5311 | -0.8988 | 0.5776 |
| Tfh | -26.8988 | -410.5044 | **-28.0473** | -18.3484 | **22.7583** |
| TGM2 luminal cell | -1.1357 | **1.7665** | -2.1383 | -5.5970 | 3.3998 |
| TNBC-specific epithelial cell | 0.2846 | -6.9020 | -1.7023 | 3.7820 | 0.7186 |
| Treg | 4.0164 | -13.8156 | -2.0061 | -18.7055 | **9.9284** |
| Venous EC | -27.2343 | -260.8903 | -38.0431 | -12.4781 | -6.1618 |

Table S35: **Coefficients of the Cox proportional hazard model** for all bayesprism sub celltypes. Coefficients having a p-Value below 0.05 are highlighted in bold.

| celltype | All | Her2 | LumA | LumB | TNBC |
|---|---|---|---|---|---|
| Arterial EC | **0.0029** | 0.3336 | 0.0712 | **0.0306** | 0.5715 |
| Breast basal cell | 0.5419 | 0.1683 | 0.5932 | 0.3493 | 0.2689 |
| Breast cancer specific luminal cell | 0.2741 | 0.5504 | 0.0684 | 0.8461 | 0.2304 |
| Breast cancer specific proliferation luminal cell | 0.2680 | 0.4422 | 0.4948 | 0.7792 | 0.5104 |
| Capillary EC | 0.9835 | 0.7064 | 0.3868 | **0.0486** | 0.9259 |
| CD4 T | 0.0895 | **0.0084** | 0.3823 | 0.4356 | 0.4661 |
| cDC2 | 0.4872 | 0.4701 | 0.9741 | 0.1168 | 0.3535 |
| CFD fibroblast | 0.7292 | 0.4267 | 0.3273 | 0.9110 | 0.3183 |
| CXCL1/2/3 fibroblast | 0.1784 | **0.0212** | 0.9576 | 0.2719 | 0.6235 |
| CXCL13 exhausted CD8 T | **0.0044** | 0.2192 | 0.2404 | 0.2046 | **0.0275** |
| GZMH CD8 T | 0.1185 | 0.3006 | 0.1844 | 0.8681 | 0.0814 |
| GZMK CD8 T | 0.4617 | 0.6509 | 0.9376 | 0.0521 | 0.5139 |
| IgA plasma | 0.6757 | **0.0175** | 0.8250 | 0.3331 | 0.4658 |
| IgG plasma cell | 0.0643 | **0.0179** | 0.1977 | 0.8263 | 0.1965 |
| INF responsed T | 0.4629 | 0.2893 | 0.7021 | 0.7717 | 0.5410 |
| Luminal progenitor | 0.4763 | 0.0801 | 0.3770 | 0.2203 | 0.7649 |
| Lymphatic EC | 0.8214 | 0.5054 | 0.2375 | 0.1433 | 0.5403 |
| Macrophage | 0.2691 | 0.4707 | 0.1653 | 0.6555 | 0.1062 |
| Mast cell | 0.8139 | 0.4482 | 0.2337 | **0.0009** | 0.3527 |
| Monocyte | **0.0238** | 0.3585 | 0.8823 | 0.0364 | 0.2143 |
| mregDC | 0.1298 | **0.0189** | 0.1699 | 0.5750 | **0.0240** |
| NK cell | 0.0631 | 0.5745 | 0.0548 | 0.5503 | 0.4839 |
| Other B cells | 0.8612 | 0.4538 | 0.1313 | 0.9141 | 0.4124 |
| Other fibroblasts | 0.1675 | 0.1260 | 0.3846 | 0.0137 | 0.5880 |
| pDC | 0.9170 | 0.0656 | 0.9113 | 0.1102 | **0.0001** |
| Pericyte | 0.2905 | **0.0063** | **3.54e-17** | **1.39e-10** | 0.3103 |
| Proliferation macrophage | 0.0955 | 0.9020 | 0.6216 | **0.0117** | **0.0208** |
| Proliferation T/NK | 0.2602 | 0.3242 | 0.6601 | 0.2851 | 0.6005 |
| Smooth muscle cell | 0.3048 | 0.2641 | 0.1356 | 0.9303 | 0.9196 |
| Tfh | 0.0388 | 0.1696 | **0.0301** | 0.6884 | **0.0019** |
| TGM2 luminal cell | 0.4735 | **0.0110** | 0.3295 | 0.1404 | 0.5081 |
| TNBC-specific epithelial cell | 0.8662 | 0.2843 | 0.6119 | 0.0617 | 0.7235 |
| Treg | 0.4820 | 0.3455 | 0.9030 | 0.1762 | **0.0257** |
| Venous EC | 0.0981 | 0.0800 | 0.2575 | 0.4400 | 0.8208 |

Table S36: **p-values of the Cox proportional hazard model** for all bayesprism sub celltypes. p-Values below 0.05 are highlighted in bold.

### 5 TCGA CybersortX

| celltype | All | Her2 | LumA | LumB | TNBC |
|---|---|---|---|---|---|
| B cell | -30.5645 | **-146.5029** | -69.2071 | -0.5262 | -26.7897 |
| Endothelial cell | -2.6442 | 1.6921 | -3.4979 | -4.3697 | -1.6703 |
| Epithelial cell | 1.0285 | **4.5129** | 0.4756 | 0.6439 | 1.4728 |
| Fibroblast | 1.6019 | **-15.4705** | 1.4337 | **7.3956** | 0.7297 |
| Myeloid cell | -1.5213 | **-11.9837** | -0.1064 | 0.4051 | -1.5463 |
| NK cell | 0.6909 | -8.2995 | 1.0484 | 0.1390 | 1.7483 |
| Perivascular cell | 2.8731 | 6.6161 | **6.8196** | 0.8350 | -2.7251 |
| Proliferation T/NK | -0.9843 | -3.7996 | 2.3543 | 2.1615 | -4.3189 |
| T cell | **-1.1227** | 0.1931 | -1.4955 | -2.1026 | 0.0227 |

Table S37: **Coefficients of the Cox proportional hazard model** for all CybersortX major celltypes using the independent level approach. Coefficients having a p-Value below 0.05 are highlighted in bold.

| celltype | All | Her2 | LumA | LumB | TNBC |
|---|---|---|---|---|---|
| B cell | 0.0995 | **0.0232** | 0.1910 | 0.9819 | 0.2085 |
| Endothelial cell | 0.1517 | 0.7382 | 0.2851 | 0.1966 | 0.6445 |
| Epithelial cell | 0.0809 | **0.0094** | 0.6289 | 0.4734 | 0.2585 |
| Fibroblast | 0.4700 | **0.0395** | 0.6477 | **0.0126** | 0.9044 |
| Myeloid cell | 0.3591 | **0.0093** | 0.9717 | 0.8962 | 0.5512 |
| NK cell | 0.4777 | 0.1249 | 0.2863 | 0.9519 | 0.3077 |
| Perivascular cell | 0.0581 | 0.1201 | **0.0153** | 0.8122 | 0.3433 |
| Proliferation T/NK | 0.4707 | 0.2271 | 0.5752 | 0.1639 | 0.0956 |
| T cell | **0.0387** | 0.9082 | 0.0519 | 0.0639 | 0.9861 |

Table S38: **p-values of the Cox proportional hazard model** for all CybersortX major using the independent level approach. p-Value below 0.05 are highlighted in bold.

| celltype | All | Her2 | LumA | LumB | TNBC |
|---|---|---|---|---|---|
| B cell | 1.0080 | -2.0793 | 5.1925 | 0.1355 | -2.1486 |
| Endothelial cell | -0.0855 | 4.6375 | -0.7636 | -0.2855 | -1.4472 |
| Epithelial cell | **1.8676** | **9.0811** | **2.6149** | -1.0672 | 0.5206 |
| Fibroblast | 1.7907 | -8.1620 | 0.5728 | 6.1803 | 6.2750 |
| Myeloid cell | -0.2141 | -2.8775 | 1.0888 | -2.2058 | 0.8939 |
| NK cell | -3.8545 | 3.6064 | -7.8695 | 1.8717 | -9.0168 |
| Perivascular cell | 4.5343 | 16.9347 | 6.8825 | 5.9400 | -1.2718 |
| Proliferation T/NK | -1.1541 | -16.4496 | 0.5130 | 6.9417 | -3.8825 |
| T cell | **-2.0098** | -4.7869 | **-3.1352** | 0.3225 | -0.1816 |

Table S39: **Coefficients of the Cox proportional hazard model** for all CybersortX major celltypes using the summing approach. Coefficients having a p-Value below 0.05 are highlighted in bold.

| celltype | All | Her2 | LumA | LumB | TNBC |
|---|---|---|---|---|---|
| B cell | 0.5844 | 0.8292 | 0.1166 | 0.9632 | 0.5809 |
| Endothelial cell | 0.9710 | 0.4251 | 0.8425 | 0.9482 | 0.7853 |
| Epithelial cell | **0.0119** | **0.0062** | **0.0063** | 0.4812 | 0.8021 |
| Fibroblast | 0.3978 | 0.2555 | 0.8413 | 0.1571 | 0.2594 |
| Myeloid cell | 0.8841 | 0.3925 | 0.6458 | 0.4855 | 0.7607 |
| NK cell | 0.3233 | 0.7464 | 0.1184 | 0.5226 | 0.4657 |
| Perivascular cell | 0.3724 | 0.3310 | 0.4569 | 0.6431 | 0.8615 |
| Proliferation T/NK | 0.7165 | 0.0536 | 0.9312 | 0.1564 | 0.5568 |
| T cell | **0.0026** | 0.1652 | **0.0004** | 0.7976 | 0.9257 |

Table S40: **p-values of the Cox proportional hazard model** for all CybersortX major celltypes using the summing approach. p-Values below 0.05 are highlighted in bold.

| celltype | All | Her2 | LumA | LumB | TNBC |
|---|---|---|---|---|---|
| B cell | -32.2873 | **-189.8166** | -93.0329 | 5.6408 | -24.5717 |
| CD4 T cell | -1.0070 | 0.3066 | **-2.5996** | 0.6155 | 0.2983 |
| CD8 T cell | -1.3177 | -4.5065 | 0.5466 | -2.2159 | -1.9327 |
| Dendritic cell | 0.5448 | 1.0380 | **3.8735** | -3.9657 | -2.8631 |
| Endothelial cell | -1.4733 | 2.6262 | -3.8111 | -2.0302 | 0.1074 |
| Epithelial cell | 1.0693 | **4.6741** | 0.3027 | 0.9482 | 1.6790 |
| Fibroblast | 1.3533 | **-17.5426** | 0.7355 | **7.7326** | 4.2966 |
| Granulocyte | 1.5295 | 1.4963 | **2.8595** | -3.1937 | 2.4779 |
| Macrophage | 1.4289 | 1.0777 | 2.5402 | 3.3707 | -2.1966 |
| Monocyte | -1.2728 | -2.2530 | -6.8876 | -1.3487 | 1.7960 |
| NK cell | 0.8231 | -15.2769 | 1.6350 | -1.0131 | 2.8166 |
| Perivascular cell | 4.1272 | 9.2994 | **8.9826** | 3.5901 | -3.4788 |
| Proliferation T/NK | -0.0126 | -2.8164 | 3.3996 | **5.2199** | -5.1262 |

Table S41: **Coefficients of the Cox proportional hazard model** for all CybersortX minor celltypes using the independent level approach. Coefficients having a p-Value below 0.05 are highlighted in bold.

| celltype | All | Her2 | LumA | LumB | TNBC |
|---|---|---|---|---|---|
| B cell | 0.1088 | **0.0412** | 0.2728 | 0.8445 | 0.2671 |
| CD4 T cell | 0.0512 | 0.8381 | **0.0008** | 0.6018 | 0.7846 |
| CD8 T cell | 0.1467 | 0.1065 | 0.6989 | 0.3187 | 0.1859 |
| Dendritic cell | 0.6034 | 0.7329 | **0.0076** | 0.1005 | 0.2675 |
| Endothelial cell | 0.4950 | 0.6503 | 0.3318 | 0.6012 | 0.9799 |
| Epithelial cell | 0.0992 | **0.0073** | 0.7784 | 0.3433 | 0.2646 |
| Fibroblast | 0.5868 | **0.0262** | 0.8333 | **0.0146** | 0.5751 |
| Granulocyte | 0.0901 | 0.6197 | **0.0124** | 0.1269 | 0.3543 |
| Macrophage | 0.3792 | 0.8158 | 0.3448 | 0.3322 | 0.5061 |
| Monocyte | 0.3825 | 0.5623 | 0.0954 | 0.7077 | 0.3708 |
| NK cell | 0.5934 | 0.0901 | 0.3375 | 0.7242 | 0.4534 |
| Perivascular cell | 0.0905 | 0.2123 | **0.0336** | 0.5475 | 0.4500 |
| Proliferation T/NK | 0.9942 | 0.4278 | 0.4970 | **0.0304** | 0.1199 |

Table S42: **p-values of the Cox proportional hazard model** for all CybersortX minor celltypes using the independent level approach. p-Values below 0.05 are highlighted in bold.

| celltype | All | Her2 | LumA | LumB | TNBC |
|---|---|---|---|---|---|
| B cell | 1.0080 | -2.0793 | 5.1925 | 0.1355 | -2.1486 |
| CD4 T cell | **-1.6920** | -0.9971 | **-3.0518** | 0.5386 | 1.0174 |
| CD8 T cell | -2.1995 | -6.3099 | -4.9113 | 0.1535 | -1.1925 |
| Dendritic cell | 0.6555 | -0.7076 | **5.4189** | -4.9126 | -5.3231 |
| Endothelial cell | -0.0855 | 4.6375 | -0.7636 | -0.2855 | -1.4472 |
| Epithelial cell | **1.8676** | **9.0811** | **2.6149** | -1.0672 | 0.5206 |
| Fibroblast | 1.7907 | -8.1620 | 0.5728 | 6.1803 | 6.2750 |
| Granulocyte | 0.1916 | 3.2861 | 4.9065 | **-19.8203** | 14.1856 |
| Macrophage | 1.5205 | -2.6807 | -3.4299 | 7.8862 | 6.3007 |
| Monocyte | -6.1145 | -7.9232 | -17.0104 | 5.5343 | -0.7546 |
| NK cell | -3.8545 | 3.6064 | -7.8695 | 1.8717 | -9.0168 |
| Perivascular cell | 4.5343 | 16.9347 | 6.8825 | 5.9400 | -1.2718 |
| Proliferation T/NK | -1.1541 | -16.4496 | 0.5130 | 6.9417 | -3.8825 |

Table S43: **Coefficients of the Cox proportional hazard model** for all CybersortX minor celltypes using the summing approach. Coefficients having a p-Value below 0.05 are highlighted in bold.

| celltype | All | Her2 | LumA | LumB | TNBC |
|---|---|---|---|---|---|
| B cell | 0.5844 | 0.8292 | 0.1166 | 0.9632 | 0.5809 |
| CD4 T cell | **0.0353** | 0.7088 | **0.0022** | 0.7984 | 0.6559 |
| CD8 T cell | 0.0718 | 0.1099 | 0.0912 | 0.9294 | 0.5430 |
| Dendritic cell | 0.7099 | 0.8897 | **0.0206** | 0.2330 | 0.2095 |
| Endothelial cell | 0.9710 | 0.4251 | 0.8425 | 0.9482 | 0.7853 |
| Epithelial cell | **0.0119** | **0.0062** | **0.0063** | 0.4812 | 0.8021 |
| Fibroblast | 0.3978 | 0.2555 | 0.8413 | 0.1571 | 0.2594 |
| Granulocyte | 0.9559 | 0.8028 | 0.2402 | **0.0174** | 0.1311 |
| Macrophage | 0.5117 | 0.5972 | 0.3369 | 0.1754 | 0.1537 |
| Monocyte | 0.2175 | 0.4714 | 0.1159 | 0.5193 | 0.9240 |
| NK cell | 0.3233 | 0.7464 | 0.1184 | 0.5226 | 0.4657 |
| Perivascular cell | 0.3724 | 0.3310 | 0.4569 | 0.6431 | 0.8615 |
| Proliferation T/NK | 0.7165 | 0.0536 | 0.9312 | 0.1564 | 0.5568 |

Table S44: **p-values of the Cox proportional hazard model** for all CybersortX minor celltypes using the summing approach. p-Values below 0.05 are highlighted in bold.

| celltype | All | Her2 | LumA | LumB | TNBC |
|---|---|---|---|---|---|
| Arterial EC | -1.8795 | 4.8244 | **-11.3987** | 4.5721 | 1.4638 |
| Breast basal cell | 12.3984 | -23.7961 | **29.9111** | -20.2442 | 10.6996 |
| Breast cancer specific luminal cell | **1.7314** | 1.9369 | **3.0999** | -1.7892 | 3.7397 |
| Breast cancer specific proliferation luminal cell | -1.0762 | 1.1540 | -5.4654 | 1.0490 | -1.9234 |
| Capillary EC | 3.2846 | 0.8178 | **10.2084** | -3.8459 | -3.5956 |
| CD4 T | -1.1449 | 0.4376 | **-2.5480** | 0.5727 | 0.8523 |
| cDC2 | 0.2481 | -0.8170 | 3.4038 | -8.9386 | -2.2260 |
| CFD fibroblast | 1.6259 | -7.3083 | 0.5129 | 5.4173 | 6.2812 |
| CXCL1/2/3 fibroblast | 5.4496 | 319.1339 | -1233.0197 | -393.3546 | 23.0349 |
| CXCL13 exhausted CD8 T | **-5.4256** | -1.1223 | **-9.4474** | -1.8589 | -4.4100 |
| GZMH CD8 T | 1.0818 | -8.0858 | 5.0644 | -0.0695 | 1.8194 |
| GZMK CD8 T | 0.7263 | -3.6779 | 1.3755 | 4.1805 | -1.1439 |
| IgA plasma | -43.5743 | -636.3439 | -1.6715 | 135.3391 | -108.8096 |
| IgG plasma cell | -68.3065 | -517.2756 | -128.7192 | 87.7805 | -54.4150 |
| INF responsed T | -4.6302 | -6.0812 | -20.0574 | 6.2810 | -2.4731 |
| Luminal progenitor | -4.6509 | 2.5084 | **-26.4033** | 2.7194 | 1.1480 |
| Lymphatic EC | -2.7765 | 10.9447 | -1.9077 | -14.3284 | 0.4273 |
| Macrophage | -5.4976 | -8.6591 | -5.9630 | -2.6794 | -5.9791 |
| Mast cell | 0.1916 | 3.2861 | 4.9065 | **-19.8203** | 14.1856 |
| Monocyte | -6.1145 | -7.9232 | -17.0104 | 5.5343 | -0.7546 |
| mregDC | -2.7398 | -24.2343 | **25.9567** | 3.8258 | -31.2626 |
| NK cell | -3.8545 | 3.6064 | -7.8695 | 1.8717 | -9.0168 |
| Other B cells | 1.1911 | 0.5849 | 5.1823 | -0.0505 | -1.6809 |
| Other fibroblasts | 7.7793 | -68.2413 | 4.7564 | **22.6254** | 2.7237 |
| pDC | 3.2878 | 9.6755 | 11.3761 | -7.9035 | -11.9171 |
| Pericyte | 26.9437 | -148.0320 | 34.9562 | -1.3019 | **47.2196** |
| Proliferation macrophage | 3.6172 | 0.7859 | -1.2147 | 8.1132 | **8.3928** |
| Proliferation T/NK | -1.1541 | -16.4496 | 0.5130 | 6.9417 | -3.8825 |
| Smooth muscle cell | 2.9662 | 18.7512 | 5.3912 | 6.4947 | -4.8754 |
| Tfh | -4.1208 | -1.5094 | -10.0390 | -0.3073 | 7.5492 |
| TGM2 luminal cell | **3.5755** | 7.2149 | **5.5518** | -1.9633 | 4.5180 |
| TNBC-specific epithelial cell | 0.2150 | -2.9131 | -8.6400 | **25.0908** | -1.7894 |
| Treg | 0.7153 | -3.9913 | 5.5743 | -1.2201 | -1.4882 |
| Venous EC | **-340.1598** | **-3996.5278** | **-337.3606** | **-15122.9634** | -172.8930 |

Table S45: **Coefficients of the Cox proportional hazard model** for all CybersortX sub celltypes. Coefficients having a p-Value below 0.05 are highlighted in bold.

| celltype | All | Her2 | LumA | LumB | TNBC |
|---|---|---|---|---|---|
| Arterial EC | 0.4884 | 0.5562 | **0.0330** | 0.3160 | 0.7819 |
| Breast basal cell | 0.2988 | 0.7328 | **0.0343** | 0.5385 | 0.6593 |
| Breast cancer specific luminal cell | **0.0233** | 0.4938 | **0.0018** | 0.2015 | 0.4735 |
| Breast cancer specific proliferation luminal cell | 0.3606 | 0.6998 | 0.1289 | 0.5382 | 0.3967 |
| Capillary EC | 0.2047 | 0.9239 | **0.0068** | 0.5605 | 0.5613 |
| CD4 T | 0.0877 | 0.8183 | **0.0032** | 0.7307 | 0.6201 |
| cDC2 | 0.9180 | 0.8963 | 0.2662 | 0.3859 | 0.6368 |
| CFD fibroblast | 0.4579 | 0.3482 | 0.8587 | 0.2740 | 0.2886 |
| CXCL1/2/3 fibroblast | 0.9409 | 0.2635 | 0.1659 | 0.0516 | 0.7332 |
| CXCL13 exhausted CD8 T | **0.0005** | 0.8193 | **0.0003** | 0.4415 | 0.1013 |
| GZMH CD8 T | 0.4969 | 0.1216 | 0.1170 | 0.9823 | 0.4493 |
| GZMK CD8 T | 0.7081 | 0.5264 | 0.6606 | 0.2658 | 0.7907 |
| IgA plasma | 0.6360 | 0.1350 | 0.9908 | 0.0722 | 0.5041 |
| IgG plasma cell | 0.1895 | 0.0835 | 0.2613 | 0.1662 | 0.3893 |
| INF responsed T | 0.4990 | 0.6912 | 0.2407 | 0.5983 | 0.8149 |
| Luminal progenitor | 0.2194 | 0.6952 | **0.0239** | 0.9209 | 0.8098 |
| Lymphatic EC | 0.6161 | 0.4141 | 0.8217 | 0.2139 | 0.9722 |
| Macrophage | 0.1143 | 0.4110 | 0.3204 | 0.7003 | 0.3593 |
| Mast cell | 0.9559 | 0.8028 | 0.2402 | **0.0174** | 0.1311 |
| Monocyte | 0.2175 | 0.4714 | 0.1159 | 0.5193 | 0.9240 |
| mregDC | 0.7002 | 0.1857 | **0.0416** | 0.7590 | 0.1205 |
| NK cell | 0.3233 | 0.7464 | 0.1184 | 0.5226 | 0.4657 |
| Other B cells | 0.5103 | 0.9418 | 0.1110 | 0.9865 | 0.6578 |
| Other fibroblasts | 0.3736 | 0.0933 | 0.7346 | **0.0497** | 0.8902 |
| pDC | 0.3863 | 0.3760 | **0.0225** | 0.3394 | 0.2743 |
| Pericyte | 0.1720 | 0.1010 | 0.3239 | 0.9739 | **0.0492** |
| Proliferation macrophage | 0.0904 | 0.8588 | 0.7163 | 0.1056 | **0.0401** |
| Proliferation T/NK | 0.7165 | 0.0536 | 0.9312 | 0.1564 | 0.5568 |
| Smooth muscle cell | 0.5655 | 0.2510 | 0.5590 | 0.6313 | 0.5348 |
| Tfh | 0.5250 | 0.8774 | 0.4648 | 0.9759 | 0.6981 |
| TGM2 luminal cell | **0.0073** | 0.0507 | **0.0019** | 0.3905 | 0.1619 |
| TNBC-specific epithelial cell | 0.9398 | 0.6960 | 0.2683 | **0.0015** | 0.6349 |
| Treg | 0.7038 | 0.4778 | 0.0611 | 0.7550 | 0.6987 |
| Venous EC | **0.0012** | **3.05e-78** | **0.0119** | **3.85e-44** | 0.1374 |

Table S46: **p-values of the Cox proportional hazard model** for all CybersortX sub celltypes. p-Value below 0.05 are highlighted in bold.

### 6 TCGA Music

| celltype | All | Her2 | LumA | LumB | TNBC |
|---|---|---|---|---|---|
| B cell | -15.0262 | **-63.7101** | -22.6750 | -1.0627 | -13.9066 |
| Endothelial cell | **1.9767** | 2.3415 | **4.3318** | 0.4540 | -0.9170 |
| Epithelial cell | 0.6277 | **3.0902** | -0.2800 | 0.8966 | 0.9763 |
| Fibroblast | 1.2098 | -13.1964 | -0.2189 | **8.0090** | 3.8823 |
| Myeloid cell | -1.5676 | -11.1384 | -2.2578 | 0.6989 | -0.3499 |
| NK cell | 0.2327 | -3.4043 | **2.9858** | **-6.0185** | 0.9393 |
| Perivascular cell | -0.6466 | -16.5175 | -0.8870 | **14.3707** | -6.8713 |
| Proliferation T/NK | -1.0868 | -0.3015 | -5.1070 | 4.2334 | -4.5293 |
| T cell | **-1.0991** | -1.5280 | **-1.8116** | -0.9978 | 0.1417 |

Table S47: **Coefficients of the Cox proportional hazard model** for all Music major celltypes using the independent level approach. Coefficients having a p-Value below 0.05 are highlighted in bold.

| celltype | All | Her2 | LumA | LumB | TNBC |
|---|---|---|---|---|---|
| B cell | 0.0888 | **0.0369** | 0.4156 | 0.9356 | 0.1489 |
| Endothelial cell | **0.0348** | 0.3618 | **0.0031** | 0.8543 | 0.6503 |
| Epithelial cell | 0.2414 | **0.0079** | 0.7553 | 0.3210 | 0.3944 |
| Fibroblast | 0.6291 | 0.1174 | 0.9538 | **0.0322** | 0.4613 |
| Myeloid cell | 0.3864 | 0.0506 | 0.5172 | 0.8478 | 0.8930 |
| NK cell | 0.8600 | 0.4966 | **0.0223** | **0.0360** | 0.8201 |
| Perivascular cell | 0.8465 | 0.3967 | 0.8728 | **0.0460** | 0.1792 |
| Proliferation T/NK | 0.5484 | 0.9313 | 0.5171 | 0.1671 | 0.1405 |
| T cell | **0.0384** | 0.3432 | **0.0358** | 0.3463 | 0.8824 |

Table S48: **p-values of the Cox proportional hazard model** for all Music major using the independent level approach. p-Values below 0.05 are highlighted in bold.

| celltype | All | Her2 | LumA | LumB | TNBC |
|---|---|---|---|---|---|
| B cell | **-12.5895** | -16.1119 | -25.1186 | -2.9773 | -19.5719 |
| Endothelial cell | **2.3719** | 3.2944 | **5.8514** | 0.8567 | -1.3676 |
| Epithelial cell | 0.8607 | **4.4313** | 0.2812 | 0.9580 | 0.6239 |
| Fibroblast | 1.5644 | -9.5816 | 0.7515 | 5.5917 | 4.6401 |
| Myeloid cell | -2.9100 | -10.3902 | -2.4607 | -4.7918 | -0.6645 |
| NK cell | -6.1467 | -17.8020 | -11.6630 | -3.2864 | 1.8471 |
| Perivascular cell | 0.9164 | -15.2761 | 4.8041 | 10.0359 | -8.4703 |
| Proliferation T/NK | -3.0635 | -68.8427 | **48.0098** | **20.3554** | -8.0451 |
| T cell | **-1.4617** | -2.3480 | -1.9764 | -2.3099 | 0.9871 |

Table S49: **Coefficients of the Cox proportional hazard model** for all Music major celltypes using the summing approach. Coefficients having a p-Value below 0.05 are highlighted in bold.

| celltype | All | Her2 | LumA | LumB | TNBC |
|---|---|---|---|---|---|
| B cell | **0.0144** | 0.4730 | 0.0642 | 0.5225 | 0.0758 |
| Endothelial cell | **0.0460** | 0.2928 | **0.0043** | 0.7963 | 0.5292 |
| Epithelial cell | 0.1389 | **0.0019** | 0.7720 | 0.3116 | 0.6049 |
| Fibroblast | 0.4380 | 0.1682 | 0.7873 | 0.1227 | 0.2909 |
| Myeloid cell | 0.0646 | 0.0502 | 0.3058 | 0.1810 | 0.8030 |
| NK cell | 0.3874 | 0.4007 | 0.4262 | 0.8355 | 0.8505 |
| Perivascular cell | 0.8233 | 0.3295 | 0.5356 | 0.1758 | 0.1912 |
| Proliferation T/NK | 0.5410 | 0.0819 | **0.0017** | **0.0282** | 0.2274 |
| T cell | **0.0365** | 0.1958 | 0.0953 | 0.0856 | 0.4615 |

Table S50: **p-values of the Cox proportional hazard model** for all Music major using the summing approach. p-Values below 0.05 are highlighted in bold.

| celltype | All | Her2 | LumA | LumB | TNBC |
|---|---|---|---|---|---|
| B cell | -16.0772 | **-61.3916** | -26.9755 | -1.4038 | -14.7935 |
| CD4 T cell | -0.6139 | 0.5235 | **-2.2186** | 1.6237 | 0.1060 |
| CD8 T cell | -1.8955 | **-6.9536** | 0.6447 | **-5.9355** | -0.0883 |
| Dendritic cell | **-7.4797** | -3.7065 | -6.2841 | -13.7109 | -7.9955 |
| Endothelial cell | **2.0398** | 2.4965 | **4.6267** | 0.7871 | -1.6314 |
| Epithelial cell | 0.6110 | **3.1671** | -0.3956 | 1.0695 | 0.9592 |
| Fibroblast | 1.2143 | -13.9620 | -0.3527 | **8.0531** | 3.6913 |
| Granulocyte | 1.3877 | 7.1218 | 1.6861 | -3.5309 | **4.8230** |
| Macrophage | 1.1738 | -13.7834 | -0.3240 | 7.0647 | 3.2809 |
| Monocyte | -2.4943 | -4.8490 | -10.0256 | 1.1635 | -1.8016 |
| NK cell | 0.1886 | -2.7974 | **7.3922** | -11.1561 | 2.2610 |
| Perivascular cell | -1.0837 | -18.3956 | -2.2306 | **16.4095** | -4.9615 |
| Proliferation T/NK | -0.7848 | 0.1810 | -7.3386 | 5.5795 | -4.2387 |

Table S51: **Coefficients of the Cox proportional hazard model** for all Music minor celltypes using the independent level approach. Coefficients having a p-Value below 0.05 are highlighted in bold.

| celltype | All | Her2 | LumA | LumB | TNBC |
|---|---|---|---|---|---|
| B cell | 0.0765 | **0.0441** | 0.3793 | 0.9171 | 0.1355 |
| CD4 T cell | 0.3472 | 0.7914 | **0.0279** | 0.2401 | 0.9291 |
| CD8 T cell | 0.0703 | **0.0154** | 0.6554 | **0.0099** | 0.9632 |
| Dendritic cell | **0.0039** | 0.6178 | 0.0962 | 0.0899 | 0.0777 |
| Endothelial cell | **0.0372** | 0.3354 | **0.0033** | 0.7549 | 0.3900 |
| Epithelial cell | 0.2502 | **0.0059** | 0.6532 | 0.2483 | 0.3980 |
| Fibroblast | 0.6237 | 0.1058 | 0.9252 | **0.0280** | 0.4646 |
| Granulocyte | 0.2238 | 0.3904 | 0.1585 | 0.3119 | **0.0047** |
| Macrophage | 0.6550 | 0.1591 | 0.9511 | 0.2716 | 0.3026 |
| Monocyte | 0.4032 | 0.5192 | 0.2728 | 0.8331 | 0.6403 |
| NK cell | 0.9471 | 0.7431 | **0.0255** | 0.0534 | 0.7560 |
| Perivascular cell | 0.7690 | 0.3023 | 0.7288 | **0.0376** | 0.3821 |
| Proliferation T/NK | 0.6820 | 0.9595 | 0.4286 | 0.1030 | 0.1972 |

Table S52: **p-values of the Cox proportional hazard model** for all Music minor celltypes using the independent level approach. p-Values below 0.05 are highlighted in bold.

| celltype | All | Her2 | LumA | LumB | TNBC |
|---|---|---|---|---|---|
| B cell | **-12.5895** | -16.1119 | -25.1186 | -2.9773 | -19.5719 |
| CD4 T cell | -0.5987 | -0.2908 | **-2.3221** | 0.8177 | 2.2329 |
| CD8 T cell | -1.1408 | -3.2134 | 2.2584 | **-4.4874** | -2.0370 |
| Dendritic cell | -3.8439 | -2.4944 | -2.8749 | -7.1366 | -3.5879 |
| Endothelial cell | **2.3719** | 3.2944 | **5.8514** | 0.8567 | -1.3676 |
| Epithelial cell | 0.8607 | **4.4313** | 0.2812 | 0.9580 | 0.6239 |
| Fibroblast | 1.5644 | -9.5816 | 0.7515 | 5.5917 | 4.6401 |
| Granulocyte | -0.2175 | 1.1490 | -1.3997 | -5.1487 | **16.5062** |
| Macrophage | 3.0362 | -10.1366 | 1.2223 | 6.8377 | 6.6964 |
| Monocyte | **-9.4069** | -12.5804 | -4.9298 | -9.4491 | -9.3925 |
| NK cell | -6.1467 | -17.8020 | -11.6630 | -3.2864 | 1.8471 |
| Perivascular cell | 0.9164 | -15.2761 | 4.8041 | 10.0359 | -8.4703 |
| Proliferation T/NK | -3.0635 | -68.8427 | **48.0098** | **20.3554** | -8.0451 |

Table S53: **Coefficients of the Cox proportional hazard model** for all Music minor celltypes using the summing approach. Coefficients having a p-Value below 0.05 are highlighted in bold.

| celltype | All | Her2 | LumA | LumB | TNBC |
|---|---|---|---|---|---|
| B cell | **0.0144** | 0.4730 | 0.0642 | 0.5225 | 0.0758 |
| CD4 T cell | 0.3480 | 0.8603 | **0.0166** | 0.5014 | 0.1202 |
| CD8 T cell | 0.2655 | 0.2826 | 0.1091 | **0.0171** | 0.3337 |
| Dendritic cell | 0.0997 | 0.7242 | 0.3909 | 0.1614 | 0.4616 |
| Endothelial cell | **0.0460** | 0.2928 | **0.0043** | 0.7963 | 0.5292 |
| Epithelial cell | 0.1389 | **0.0019** | 0.7720 | 0.3116 | 0.6049 |
| Fibroblast | 0.4380 | 0.1682 | 0.7873 | 0.1227 | 0.2909 |
| Granulocyte | 0.9303 | 0.9743 | 0.5860 | 0.5081 | **0.0006** |
| Macrophage | 0.2898 | 0.2207 | 0.8081 | 0.3069 | 0.1437 |
| Monocyte | **0.0308** | 0.3181 | 0.7094 | 0.3937 | 0.0843 |
| NK cell | 0.3874 | 0.4007 | 0.4262 | 0.8355 | 0.8505 |
| Perivascular cell | 0.8233 | 0.3295 | 0.5356 | 0.1758 | 0.1912 |
| Proliferation T/NK | 0.5410 | 0.0819 | **0.0017** | **0.0282** | 0.2274 |

Table S54: **p-values of the Cox proportional hazard model** for all Music minor celltypes using the summing approach. p-Values below 0.05 are highlighted in bold.

| celltype | All | Her2 | LumA | LumB | TNBC |
|---|---|---|---|---|---|
| Arterial EC | 4.1480 | 5.7588 | 5.6792 | 5.9561 | 0.2513 |
| Breast basal cell | -2.3258 | -5.1388 | -5.7774 | 9.7724 | 0.1604 |
| Breast cancer specific luminal cell | 0.4963 | 0.4725 | 0.2547 | -0.1758 | **4.8301** |
| Breast cancer specific proliferation luminal cell | 0.5299 | 3.1262 | -2.8391 | 3.0240 | -0.4823 |
| Capillary EC | 2.7170 | -1.6411 | **8.3402** | -2.0976 | -3.3801 |
| CD4 T | -0.5431 | -0.4632 | **-1.8841** | 0.2566 | 1.9773 |
| cDC2 | -6.1000 | -3.0669 | -8.3753 | -7.8065 | -3.5097 |
| CFD fibroblast | 0.9607 | -9.8444 | 0.7959 | 5.7762 | 1.2315 |
| CXCL1/2/3 fibroblast | **14.4911** | -2770.3697 | -57.4716 | -45.1035 | **16.0235** |
| CXCL13 exhausted CD8 T | **-7.9459** | -7.8050 | -81.9326 | -1.1202 | -9.0308 |
| GZMH CD8 T | -7.9645 | -13.3483 | -1744.1759 | -14.3721 | 19.1266 |
| GZMK CD8 T | -0.4547 | -2.2583 | **2.8079** | **-4.7062** | -0.5602 |
| IgA plasma | -2.6353 | -682.5341 | 8.8293 | **38.5866** | -126.9289 |
| IgG plasma cell | **-49.2270** | **-201.9568** | -270.3538 | -17.4386 | -31.3686 |
| INF responsed T | -11.5394 | -73.3893 | 12.3829 | -15.9049 | -18.1087 |
| Luminal progenitor | -2.6672 | 1.2691 | **-13.0901** | -7.3820 | -0.8265 |
| Lymphatic EC | 3.1791 | 26.6210 | **5.9853** | -1.8252 | 1.0355 |
| Macrophage | 0.4123 | -9.4764 | -3.4115 | 4.1899 | 4.9643 |
| Mast cell | -0.2175 | 1.1490 | -1.3997 | -5.1487 | **16.5062** |
| Monocyte | **-9.4069** | -12.5804 | -4.9298 | -9.4491 | -9.3925 |
| mregDC | -0.4682 | -117.1112 | **49.8565** | -16.2705 | -25.1282 |
| NK cell | -6.1467 | -17.8020 | -11.6630 | -3.2864 | 1.8471 |
| Other B cells | **-10.7167** | -6.4657 | -22.9321 | -3.9598 | -16.5354 |
| Other fibroblasts | 4.3754 | -25.0279 | 4.5560 | **25.7106** | **-1446.6078** |
| pDC | -0.5184 | 2.6703 | 3.5524 | -6.3674 | -2.9030 |
| Pericyte | 0.9251 | -143.1812 | -2.5358 | **32.5243** | -41.7658 |
| Proliferation macrophage | 8.4515 | -5.2617 | 20.6023 | 10.6069 | 8.7878 |
| Proliferation T/NK | -3.0635 | -68.8427 | **48.0098** | **20.3554** | -8.0451 |
| Smooth muscle cell | 0.9353 | -7.3686 | 5.5805 | 5.1091 | -5.5453 |
| Tfh | -8.4608 | 13.6374 | -14.1236 | -19.0414 | -15.0472 |
| TGM2 luminal cell | **1.9472** | **6.5208** | **2.4496** | 0.1378 | -0.7194 |
| TNBC-specific epithelial cell | 0.7026 | 17.1293 | NaN | NaN | 0.2678 |
| Treg | **6.4156** | 4.5115 | 3.9496 | **10.2100** | 1.5692 |
| Venous EC | -3.1182 | -3.7308 | -1.8189 | 1.4798 | -10.6539 |

Table S55: **Coefficients of the Cox proportional hazard model** for all Music sub celltypes. Coefficients having a p-Value below 0.05 are highlighted in bold.

| celltype | All | Her2 | LumA | LumB | TNBC |
|---|---|---|---|---|---|
| Arterial EC | 0.0536 | 0.1743 | 0.0935 | 0.3963 | 0.9486 |
| Breast basal cell | 0.3481 | 0.7767 | 0.1572 | 0.5831 | 0.9617 |
| Breast cancer specific luminal cell | 0.4334 | 0.7861 | 0.7888 | 0.8398 | **0.0010** |
| Breast cancer specific proliferation luminal cell | 0.5934 | 0.3374 | 0.2442 | 0.0779 | 0.7713 |
| Capillary EC | 0.1513 | 0.7683 | **0.0011** | 0.6290 | 0.4239 |
| CD4 T | 0.3511 | 0.7705 | **0.0295** | 0.8307 | 0.1237 |
| cDC2 | 0.0782 | 0.7065 | 0.1129 | 0.4432 | 0.5724 |
| CFD fibroblast | 0.6723 | 0.2381 | 0.7851 | 0.2427 | 0.8242 |
| CXCL1/2/3 fibroblast | **4.00e-5** | 0.3142 | 0.3647 | 0.3786 | **3.12e-13** |
| CXCL13 exhausted CD8 T | **0.0301** | 0.3710 | 0.0517 | 0.8506 | 0.1047 |
| GZMH CD8 T | 0.3616 | 0.3908 | 0.2612 | 0.4398 | 0.3088 |
| GZMK CD8 T | 0.6879 | 0.5224 | **0.0428** | **0.0115** | 0.8540 |
| IgA plasma | 0.9521 | 0.0913 | 0.8950 | **7.00e-5** | 0.2221 |
| IgG plasma cell | **0.0372** | **0.0319** | 0.0621 | 0.6270 | 0.1707 |
| INF responsed T | 0.1583 | 0.1636 | 0.3398 | 0.2773 | 0.3050 |
| Luminal progenitor | 0.1226 | 0.6494 | **0.0278** | 0.2075 | 0.7029 |
| Lymphatic EC | 0.1356 | 0.0646 | **0.0461** | 0.7770 | 0.7703 |
| Macrophage | 0.8924 | 0.3540 | 0.5759 | 0.5393 | 0.2721 |
| Mast cell | 0.9303 | 0.9743 | 0.5860 | 0.5081 | **0.0006** |
| Monocyte | **0.0308** | 0.3181 | 0.7094 | 0.3937 | 0.0843 |
| mregDC | 0.9780 | 0.0650 | **0.0025** | 0.4981 | 0.5158 |
| NK cell | 0.3874 | 0.4007 | 0.4262 | 0.8355 | 0.8505 |
| Other B cells | **0.0333** | 0.7130 | 0.0599 | 0.4162 | 0.0998 |
| Other fibroblasts | 0.4675 | 0.4468 | 0.6278 | **0.0272** | **1.86e-05** |
| pDC | 0.8878 | 0.8065 | 0.5257 | 0.3719 | 0.7480 |
| Pericyte | 0.9314 | 0.1635 | 0.9286 | **0.0122** | 0.0559 |
| Proliferation macrophage | 0.1191 | 0.6952 | 0.0733 | 0.3766 | 0.3300 |
| Proliferation T/NK | 0.5410 | 0.0819 | **0.0017** | **0.0282** | 0.2274 |
| Smooth muscle cell | 0.8360 | 0.5979 | 0.5088 | 0.5752 | 0.4228 |
| Tfh | 0.2064 | 0.2488 | 0.1293 | 0.4386 | 0.4510 |
| TGM2 luminal cell | **0.0096** | **0.0362** | **0.0061** | 0.9271 | 0.8862 |
| TNBC-specific epithelial cell | 0.8335 | 0.1608 | NaN | NaN | 0.9384 |
| Treg | **0.0201** | 0.5308 | 0.4446 | **0.0110** | 0.7881 |
| Venous EC | 0.3797 | 0.6876 | 0.6786 | 0.8766 | 0.3185 |

Table S56: **p-values of the Cox proportional hazard model** for all Music sub celltypes. p-Values below 0.05 are highlighted in bold.

### 7 TCGA Benchmark summary

| Method | Layer | All | Her2 | LumA | LumB | TNBC | Total |
|---|---|---|---|---|---|---|---|
| HIDE | Major | 0 | 4 | 1 | 2 | 2 | 9 |
| | Minor | 1 | 5 | 2 | 2 | 3 | 13 |
| | Sub | 4 | 6 | 4 | 7 | 6 | 27 |
| Bayesprism | Major (ILA) | 0 | 3 | 0 | 2 | 2 | 7 |
| | Major (sum. apr.) | 0 | 3 | 1 | 0 | 0 | 4 |
| | Minor (ILA) | 1 | 3 | 0 | 2 | 3 | 9 |
| | Minor (sum. apr.) | 2 | 2 | 1 | 2 | 1 | 8 |
| | Sub | 3 | 7 | 2 | 6 | 6 | 24 |
| CybersortX | Major (ILA) | 1 | 4 | 1 | 1 | 0 | 7 |
| | Major (sum. apr.) | 2 | 1 | 2 | 0 | 0 | 5 |
| | Minor (ILA) | 0 | 3 | 4 | 2 | 0 | 9 |
| | Minor (sum. apr.) | 2 | 1 | 3 | 1 | 0 | 7 |
| | Sub | 4 | 1 | 11 | 4 | 2 | 22 |
| Music | Major (ILA) | 2 | 2 | 3 | 3 | 0 | 10 |
| | Major (sum. apr.) | 3 | 1 | 2 | 1 | 0 | 7 |
| | Minor (ILA) | 2 | 3 | 3 | 3 | 1 | 12 |
| | Minor (sum. apr.) | 3 | 1 | 3 | 2 | 1 | 10 |
| | Sub | 7 | 2 | 8 | 6 | 4 | 27 |

Table S57: **Summary of significant p-Values of the TCGA-BRCA survival analysis per layer.** "Sum. apr." refers to the summing approach, "ILA" to the individual level approach.

| | BayesPrism | | CIBERSORTx | | MuSiC | | HIDE |
|---|---|---|---|---|---|---|---|
| | sum. apr. | ILA | sum. apr. | ILA | sum. apr. | ILA | |
| inconsistent | 0 | 3 | 0 | 3 | 0 | 7 | 0 |
| consistent | 4 | 4 | 4 | 5 | 10 | 7 | 15 |

Table S58: **Number of consistensies and inconsistensies between the cell type levels for the cell types significant for survival.** All cell types which are identical between major, minor and sub-minor are regarded or cell types which just have one child term. We count an inconsistency when a cell type is significant in one level but the same cell type is not significant in another level or has a different sign. Cell types are counted consistent when they are significant in two levela and have the same sign.

| | Bayesprism | CybersortX | Music | HIDE |
|---|---|---|---|---|
| major level | 4 | 1 | 6 | 5 |
| minor level | 2 | 2 | 2 | 6 |
| sub level | 7 | 3 | 8 | 11 |
| total | 13 | 6 | 16 | 22 |

Table S59: Count of significant celltypes across the breast cancer subtypes linked in a chain. The major and minor levels of Bayesprism, CybersortX and Music have been estimated by the independent level approach. A chain is defined by a significant celltype that has either a significant parent or child celltype. In the case of a celltype having only one major and various subtypes, but no minor celltype, the chain is defined as a significant in a parent cell type in at least one of its corresponding child cell types.

| | Bayesprism | CybersortX | Music | HIDE |
|---|---|---|---|---|
| major level | 3 | 4 | 3 | 5 |
| minor level | 2 | 3 | 2 | 6 |
| sub level | 5 | 9 | 7 | 11 |
| total | 10 | 16 | 12 | 22 |

Table S60: **Count of significant celltypes of the TCGA-BRCA survival analysis across the breast cancer subtypes linked in a chain**. The major and minor levels of Bayesprism, CybersortX and Music have been estimated by the summing approach. In the case of a celltype having only one major and various subtypes, but no minor celltype, the chain is defined as a significant celltype in a parent cell type and at in at least one of its corresponding child cell types.
